# Image-Conditioned Diffusion Preserves Latent Physiological Signals and Enables Cross-Site Transfer of Synthetic Medical Images

**DOI:** 10.64898/2026.05.04.722524

**Authors:** Doron Yaya-Stupp, Guy Lutsker, Or Spiegel-Yerushalmi, Eran Segal

**Affiliations:** Department of Computer Science and Applied Mathematics, Weizmann Institute of Science, Rehovot, Israel; The Gertner Institute for Epidemiology and Health Policy Research, Sheba Medical Center, Ramat Gan, Israel; Mohamed bin Zayed University of Artificial Intelligence, Abu Dhabi, UAE

**Keywords:** image-conditioned diffusion, synthetic medical images, biomarker preservation, privacy-utility tradeoff, cross-site transfer learning, retinal fundus imaging, chest radiography

## Abstract

Medical imaging models depend on large, shareable datasets, yet privacy constraints limit data dissemination. Text-conditioned diffusion models generate only what the prompt specifies and can miss subtle, distributed clinical signals, such as continuous physiological biomarkers, that downstream physiological models rely on. Here we systematically evaluate image-conditioned (I2I) diffusion, which anchors generation to a real source image. Fine-tuning Stable Diffusion on retinal fundus photographs from the Human Phenotype Project, we find that I2I-synthetic images retain more source-specific signal than text-conditioned generation, including for biomarkers absent from the prompt (hemoglobin, waist circumference). Estimators pretrained on I2I-synthetic images alone transfer across cohorts (Human Phenotype Project to UK Biobank) at performance comparable to real-image pretraining, most clearly when target-site labels are scarce. Finally, varying the I2I noise strength provides a tunable control on empirical re-identification risk. Under similarity-based linkage attacks, image-based linkage falls substantially with increasing strength for retinal images, whereas chest radiographs remain highly linkable at all strengths tested.

**Data and Code Availability:** This study uses three imaging cohorts under their respective terms: the Human Phenotype Project (HPP) (Reicher et al., 2025; Shilo et al., 2021) and UK Biobank (UKB) (Sudlow et al., 2015) retinal images, available to approved researchers on application, and the fully public NIH CXR-8 chest radiographs (Wang et al., 2017); per-cohort access details are in Appendix A. Code to train LoRA adapters and biomarker estimator models, as well as to reproduce the analyses, is available at https://github.com/dst1/SynthMed-demo. Trained LoRA adapters can be found on HuggingFace: https://huggingface.co/doronys/synthmed-loras.

**Institutional Review Board (IRB):** The Human Phenotype Project (HPP) cohort study is conducted according to the principles of the Declaration of Helsinki and was approved by the Institutional Review Board (IRB) of the Weizmann Institute of Science (reference number 1719-1). All HPP participants signed an informed consent form to be included in the cohort, and all identifying details of the participants were removed prior to computational analysis. Data from the UK Biobank was accessed under application ID 28784, complying with the UK Biobank’s generic ethical approval from the NHS National Research Ethics Service. The NIH CXR-8 dataset is a fully de-identified, publicly available dataset and is therefore exempt from additional IRB review.

## 1. Introduction

Medical image models require the curation and annotation of large image datasets. Recent image foundation models for medicine promise to reduce labeleddata requirements by enabling efficient fine-tuning on smaller task-specific datasets. However, privacy constraints and concerns raised about the potential reidentification of patients from images used for training may hamper their development and dissemination (Packhäuser et al., 2022; Sun et al., 2025). Synthetic images provide an alternative data source, which motivates approaches that simultaneously quantify (i) downstream clinical utility—such as enabling crossinstitutional transfer learning without sharing restricted source images—and (ii) re-identification risk.

Yet current medical image generators are mostly text- or label-conditioned and can only generate what the prompt enumerates (Section 2). In medical imaging, clinically relevant signals are often subtle, distributed, and correlated across phenotypes, and may not be captured by a small set of explicit conditioning variables. These latent physiological signals and correlated variation can thus be lost unless explicitly mentioned, especially when seldom known in advance. This limitation is particularly important for continuous biomarkers (e.g., hemoglobin level, blood pressure, etc.) whose imaging correlates may be weak and not directly interpretable, yet still measurable by learned models (Lang et al., 2024; Poplin et al., 2018).

Conditioning diffusion on both the text and a noised version of an existing image (Meng et al., 2022), henceforth referred to as image-to-image (I2I), in contrast to text-to-image (T2I), can thus portray latent attributes in the original image that were not explicitly mentioned in the generation prompt. Moreover, it offers the opportunity to adjust and quantify the tradeoff between privacy and image quality. Modulating the strength of noise injected into the source image provides a natural knob that interpolates between near-identity transformations and unconstrained generation, enabling a controllable privacy–utility spectrum.

In this paper, we systematically evaluate text- and image-conditioned diffusion to generate medical images that preserve the source-specific signal of the original image when creating a synthetic counterpart. We treat I2I generation as a stochastic transformation that produces a synthetic counterpart of a real image while preserving downstream-relevant signals and providing a tunable control on re-identification risk. We show that this approach better preserves both disease classes and source-specific attributes in retina images from the Human Phenotype Project (HPP) (Reicher et al., 2025; Shilo et al., 2021) and chest X-ray (CXR) images from the NIH CXR8 dataset (Wang et al., 2017) using several complementary metrics: (i) pixel and perceptual similarity, embedding-space proximity, and (iii) agreement of trained clinical biomarker estimators between real images and their synthetic counterparts. We further show that the resulting synthetic images can be used to train biomarker estimator models using syntheticonly data with emphasis on the low-data regime, and that synthetic-only I2I pretraining transfers to an external cohort (UK Biobank (Sudlow et al., 2015)) at performance comparable to real-image pretraining. We additionally evaluate whether mixing real and synthetic images improves performance relative to real-only training under controlled subsampling. Finally, we characterize the privacy–utility frontier as a function of the I2I noise strength *s*, providing empirical re-identification risk estimates.

Our contributions are:

1. **Signal beyond the prompt:** I2I retains more source-specific signal than T2I, including for biomarkers absent from the prompt.
2. **Synthetic-only training and cross-site transfer:** on retinal images, estimators pretrained on I2I-synthetic images alone transfer to an external cohort at performance comparable to real-image pretraining, most clearly with scarce target labels. Mixing synthetic with real images gave limited gains, mainly with small training sets.
3. **A tunable control on empirical reidentification risk:** the I2I noise strength tunes image-based linkage under similarity-based attacks, substantially reducing it for retinal images, whereas CXR remains highly linkable at all tested strengths.

## 2. Related Work

### Synthetic medical images

Generative models of medical images, and predictive models trained on their outputs, have shown promise for CXR (Bluethgen et al., 2025; Han et al., 2020), retina fundus images (Wang et al., 2025), and other modalities (Du-Mont Schütte et al., 2021; Han et al., 2018; Ktena et al., 2024). Previous work has typically focused on discrete disease labels, such as pneumonia presence (Bluethgen et al., 2025) or diabetic retinopathy grade (Sun et al., 2025; Wang et al., 2025), with generation limited to the attributes they target. We instead target source-image signal, including continuous biomarkers never named in the prompt.

### Image-conditioned diffusion

Starting the diffusion process from a noised version of an existing image, jointly with a text prompt, is an established technique in computer vision (Meng et al., 2022). We adopt it as our I2I generator and use its noise strength as a tunable control on re-identification risk; work on memorization and privacy of diffusion-generated data is discussed in Section 5.

## 3. Methods

### 3.1. Datasets

We used three imaging cohorts. The **Human Phenotype Project (HPP)** is a large-scale prospective cohort with deep clinical and molecular phenotyping (Reicher et al., 2025; Shilo et al., 2021); we used retinal fundus images from 10,010 participants (23,259 images) with clinical data recorded up to or at the fundoscopy visit—demographics, vitals, anthropometrics, and comprehensive laboratory blood panels (age, sex, blood pressure, grip strength, waist circumference, BMI, hemoglobin, HbA1c%, creatinine, and cholesterol, among others). Participants were split 90%/10% into train/test with no participant crossing splits, and 5-fold cross-validation splits were derived analogously from the training set (Table 1). The **UK Biobank (UKB)** (Sudlow et al., 2015) contributed retinal images from 17,639 participants (35,334 images; first instance, first visit), processed to match the HPP shared attribute set (age, sex, hemoglobin, blood pressure, grip strength, waist circumference, creatinine; Table 2). **NIH CXR-8** (Wang et al., 2017) is a public dataset of 108,948 frontal chest X-rays from 32,717 participants with 14 text-derived labels; we retained 12 after removing Edema and Pneumonia, kept the first PA-view image per participant, downsampled no-finding images to 30%, and split 80%/20% into train/test (Table 3). Exact attribute lists and preprocessing for all cohorts are in Appendix A.

**Table 1:** Cohort characteristics. Sample, demographic, and clinical characteristics of the three imaging cohorts, shown for the overall dataset and by train/test split. Continuous variables are reported as mean (SD); binary/categorical variables as N (%); images per person as mean (SD). Cohort composition, shared biomarkers, and label definitions are given in the text above. BP – blood pressure; Circ. – circumference; Thick. – thickening.

| **Human Phenotype Project** |  |  |  |
|  | Overall | Train | Test |
|---|---|---|---|
| # images | 23,259 | 20,917 | 2,342 |
| # unique persons | 10,010 | 9,009 | 1,001 |
| Visits per person | 1.2 (0.4) | 1.2 (0.4) | 1.2 (0.4) |
| Images per person | 2.3 (0.8) | 2.3 (0.8) | 2.3 (0.8) |
| Age | 52.6 (8.0) | 52.5 (8.0) | 53.1 (8.1) |
| Sex – female | 11,620 (52.8%) | 10,441 (52.7%) | 1,179 (53.3%) |
| Eye side – Right | 11,639 (50.0%) | 10,466 (50.0%) | 1,173 (50.1%) |
| Hemoglobin | 14.1 (1.2) | 14.1 (1.2) | 14.1 (1.2) |
| BP (Systolic) | 119.4 (16.6) | 119.3 (16.7) | 119.9 (16.2) |
| Waist Circ. | 88.4 (12.1) | 88.4 (12.0) | 88.5 (12.4) |
| Creatinine | 0.8 (0.2) | 0.8 (0.2) | 0.8 (0.2) |
| Grip | 76.5 (26.3) | 76.4 (26.3) | 76.6 (26.4) |

**Table 2:** Cohort characteristics (continued) — UK Biobank.

| **UK Biobank** |  |  |  |
|  | Overall | Train | Test |
|---|---|---|---|
| # images | 35,334 | 31,803 | 3,531 |
| # unique persons | 17,639 | 15,875 | 1,764 |
| Images per person | 2.0 (0.1) | 2.0 (0.1) | 2.0 (0.0) |
| Age | 54.3 (8.2) | 54.3 (8.2) | 54.0 (8.3) |
| Sex – female | 20,014 (56.6%) | 17,995 (56.6%) | 2,019 (57.2%) |
| Eye side – Right | 17,665 (50.0%) | 15,899 (50.0%) | 1,766 (50.0%) |
| Hemoglobin | 14.3 (1.2) | 14.3 (1.2) | 14.3 (1.2) |
| BP (Systolic) | 137.6 (19.2) | 137.6 (19.2) | 137.9 (19.3) |
| Waist Circ. | 89.0 (13.2) | 89.0 (13.1) | 89.1 (13.5) |
| Creatinine | 0.8 (0.2) | 0.8 (0.2) | 0.8 (0.2) |
| Grip | 31.3 (11.1) | 31.3 (11.1) | 31.3 (10.9) |

**Table 3:**
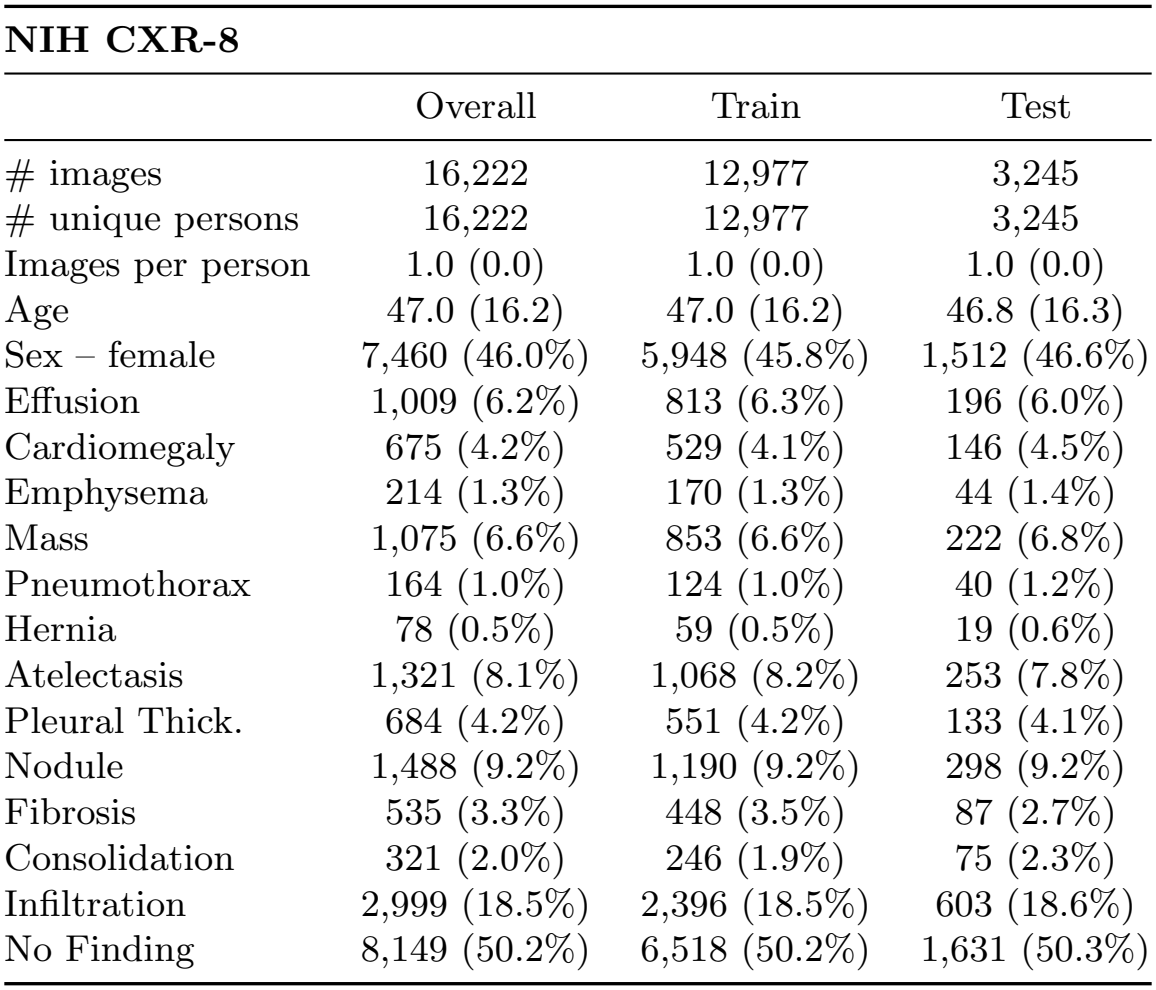
Cohort characteristics (continued) — NIH CXR-8.

| **NIH CXR-8** |  |  |  |
|  | Overall | Train | Test |
|---|---|---|---|
| # images | 16,222 | 12,977 | 3,245 |
| # unique persons | 16,222 | 12,977 | 3,245 |
| Images per person | 1.0 (0.0) | 1.0 (0.0) | 1.0 (0.0) |
| Age | 47.0 (16.2) | 47.0 (16.2) | 46.8 (16.3) |
| Sex – female | 7,460 (46.0%) | 5,948 (45.8%) | 1,512 (46.6%) |
| Effusion | 1,009 (6.2%) | 813 (6.3%) | 196 (6.0%) |
| Cardiomegaly | 675 (4.2%) | 529 (4.1%) | 146 (4.5%) |
| Emphysema | 214 (1.3%) | 170 (1.3%) | 44 (1.4%) |
| Mass | 1,075 (6.6%) | 853 (6.6%) | 222 (6.8%) |
| Pneumothorax | 164 (1.0%) | 124 (1.0%) | 40 (1.2%) |
| Hernia | 78 (0.5%) | 59 (0.5%) | 19 (0.6%) |
| Atelectasis | 1,321 (8.1%) | 1,068 (8.2%) | 253 (7.8%) |
| Pleural Thick. | 684 (4.2%) | 551 (4.2%) | 133 (4.1%) |
| Nodule | 1,488 (9.2%) | 1,190 (9.2%) | 298 (9.2%) |
| Fibrosis | 535 (3.3%) | 448 (3.5%) | 87 (2.7%) |
| Consolidation | 321 (2.0%) | 246 (1.9%) | 75 (2.3%) |
| Infiltration | 2,999 (18.5%) | 2,396 (18.5%) | 603 (18.6%) |
| No Finding | 8,149 (50.2%) | 6,518 (50.2%) | 1,631 (50.3%) |

**Table 4:** Image augmentation pipeline parameters.

|  | **Retina** | **CXR** |
|---|---|---|
| *Rotation* |  |  |
| rotation_degrees | 15 | 10 |
| rotation_p | 0.7 | 0.7 |
| *Flips* |  |  |
| horizontal_flip_p | 0 | 0 |
| vertical_flip_p | 0.1 | 0 |
| *Center Crop* |  |  |
| crop_size_factor | 1 | 0.82 |
| center_crop_p | 0 | 0.7 |
| *Color Jitter* |  |  |
| brightness | 0.1 | 0.2 |
| contrast | 0.05 | 0.15 |
| saturation | 0.05 | 0.15 |
| hue | 0.04 | 0 |
| color_jitter_p | 0.1 | 0.1 |

### 3.2. Diffusion Model and Conditioning Protocol

We fine-tuned Stable Diffusion 1.4 (Rombach et al., 2022), which supports both text and image conditioning, on the training set of each dataset using lowrank adapters (LoRA) (Hu et al., 2022) applied to the UNet (rank 64, *α* = 32) through the Hugging-Face Diffusers library (von Platen et al., 2022); the full training configuration is in Appendix B. We then generated synthetic images under two regimes: text-to-image (T2I), sampled from pure noise conditioned only on the structured prompt; and image-to-image (I2I), in which each synthetic image is a stochastic transformation of a specific real source image guided by the same structured prompt. For I2I, the source is noised up to a point in the forward schedule set by a strength parameter *s* and then denoised under the same prompt: at *s* = 0 the output is the source, higher strengths approach T2I, and lower strengths preserve more source structure. Because the DDIM schedule leaves residual source signal even at *s* = 1, T2I generations are reported separately rather than as *s* = 1. Unless otherwise specified, all I2I results use *s* = 0.5 as a reference strength for utility; it is not a release setting (Section 5).

### 3.3. Prompting

For both modalities we built prompts from a structured template encoding modality/view descriptors and the experiment’s clinical attributes (e.g., age, sex, and systolic blood pressure for retina; radio-graphic findings for CXR), or modality only for unconditioned models. T2I generations were produced from these prompts alone; I2I additionally used a real source image as the starting point. Full templates and worked examples are in Table 5.

**Table 5:**
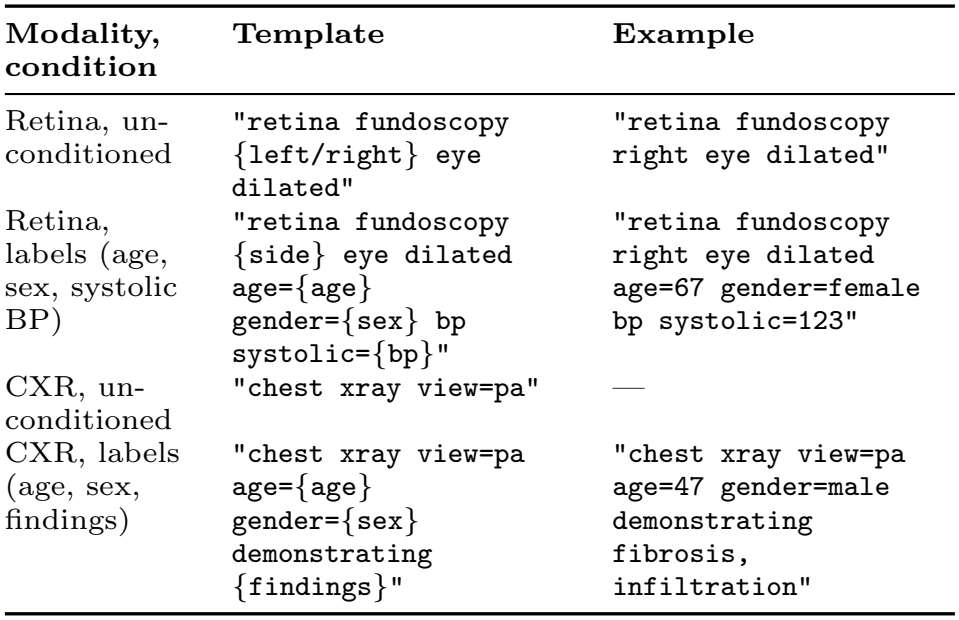
Structured prompt templates by modality and conditioning condition.

| **Modality, condition** | **Template** | **Example** |
|---|---|---|
| Retina, un-conditioned | "retina fundoscopy {left/right} eye dilated" | "retina fundoscopy right eye dilated" |
| Retina, labels (age, sex, systolic BP) | "retina fundoscopy {side} eye dilated age={age} gender={sex} bp systolic={bp}" | "retina fundoscopy right eye dilated age=67 gender=female bp systolic=123" |
| CXR, un-conditioned | "chest xray view=pa" | — |
| CXR, labels (age, sex, findings) | "chest xray view=pa age={age} gender={sex} demonstrating {findings}" | "chest xray view=pa age=47 gender=male demonstrating fibrosis, infiltration" |

### 3.4. Estimator Models

All estimator models used a ResNet v2 101 pretrained with the BiT recipe (Kolesnikov et al., 2020) from TIMM (Wightman, 2019), with per-attribute heads (cross-entropy for classification, mean-squared error for regression, missing values masked). We trained for 20,000 steps with AdamW and early stopping on validation loss (patience 10), using separate constant learning rates for the pretrained base and the heads. Learning rates and the base architecture were selected by grid search (Appendix B).

### 3.5. Training Regimes and Cross-Site Transfer

We evaluated estimator training under four regimes: real-only; real with conventional augmentations (rotation, photometric perturbations, and cropping; Figure 6); synthetic-only (T2I or I2I); and real:synthetic mixtures at predefined ratios (1X–10X synthetic per real). Synthetic images inherited their source subject’s split to prevent leakage. For the low-data regime, we additionally trained on nested subsamples, with smaller sets strict subsets of larger ones. For cross-site transfer, we pretrained estimators on HPP under each regime and fine-tuned on UKB using matched phenotype definitions and preprocessing, evaluating performance as a function of UKB training-set size. All comparisons used identical architectures, optimization settings, early-stopping criteria, and evaluation pipelines across regimes; augmentation pipeline parameters are in Appendix B.

### 3.6. Evaluation Metrics

Estimators were evaluated by Pearson correlation (regression targets) and AUROC (classification), with 95% percentile-bootstrap confidence intervals over the held-out test set. Regime comparisons (e.g., I2I vs T2I, synthetic-only vs real-only, mixed vs augmented) used paired bootstraps over shared test examples, reporting both nominal and Benjamini– Hochberg FDR-adjusted results. Image generation was evaluated with pixel-level PSNR and SSIM (Wang et al., 2004) and the neural-network metrics FID (Heusel et al., 2017) and LPIPS (VGG backbone) (Zhang et al., 2018), all computed with pyiqa 0.14 (Chen and Mo, 2022).

### 3.7. Structural and Semantic Preservation

We quantified higher-level “semantic” similarity using self-supervised DINOv2 embeddings (Oquab et al., 2024): images were embedded with a frozen encoder and compared by cosine distance (real-to-real and real-to-synthetic). These embeddings also served for UMAP visualization of real vs. synthetic manifold overlap.

### 3.8. Re-Identification Risk Evaluation

We evaluated privacy risk with similarity-based linkage attacks that match each of 300 test-set synthetic images to its nearest real image in the test set under a pixel-space measure (SSIM) and an embedding-space measure (frozen-encoder cosine distance), recording whether the true source was recovered as the top-1 match. Results were contextualized against a real-to-real “distance to closest record” baseline and repeated across I2I strengths. We also evaluated attackers that know the prompted attributes of every gallery record: an attribute-only attacker picks uniformly among records whose attributes match the synthetic image’s prompt (expected top-1 = mean inverse match-set size), and attributes + image attackers rank those records by embedding cosine (attributes + embedding) or SSIM (attributes + SSIM). All but the SSIM attacks were repeated on a gallery of 19,075 retinal and 16,222 CXR training and test images.

## 4. Results

### 4.1. Image-Conditioning Retains Source-Specific Signal Beyond the Prompt

Using the fine-tuned generator and structured prompts (Section 3.3), we compared T2I generations with I2I counterparts at the reference strength *s* = 0.5.

Generated images showed consistently greater similarity to their corresponding source images under I2I than under T2I, both in pixel space (Figure 1 – PSNR, SSIM) and in semantic latent measures (Figure 1 – LPIPS, FID). Representative fundus examples illustrate that I2I better retains clinically salient structure (e.g., vessel geometry and optic disc appearance) while allowing controlled variation, whereas T2I generations more frequently drift in global appearance (Figure 7).

**Figure 1:**
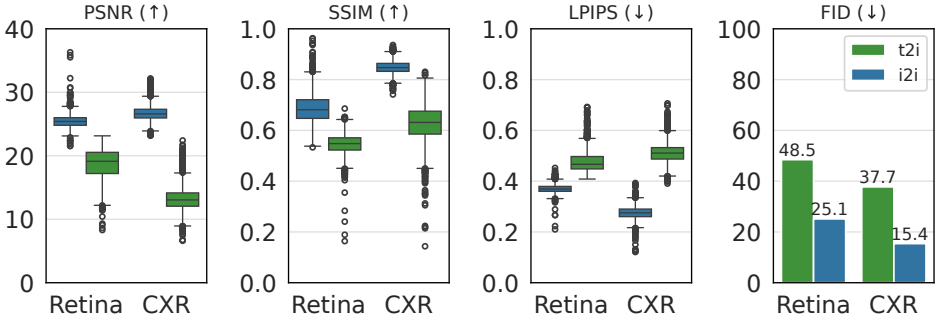
Image-to-image vs text-to-image synthesis fidelity. Image-fidelity metrics for Retina and Chest X-ray (CXR), I2I (blue) vs T2I (green): PSNR and SSIM (higher is better); LPIPS and FID (lower is better). Box plots: median, IQR, 1.5 × IQR whiskers.

**Figure 2:**
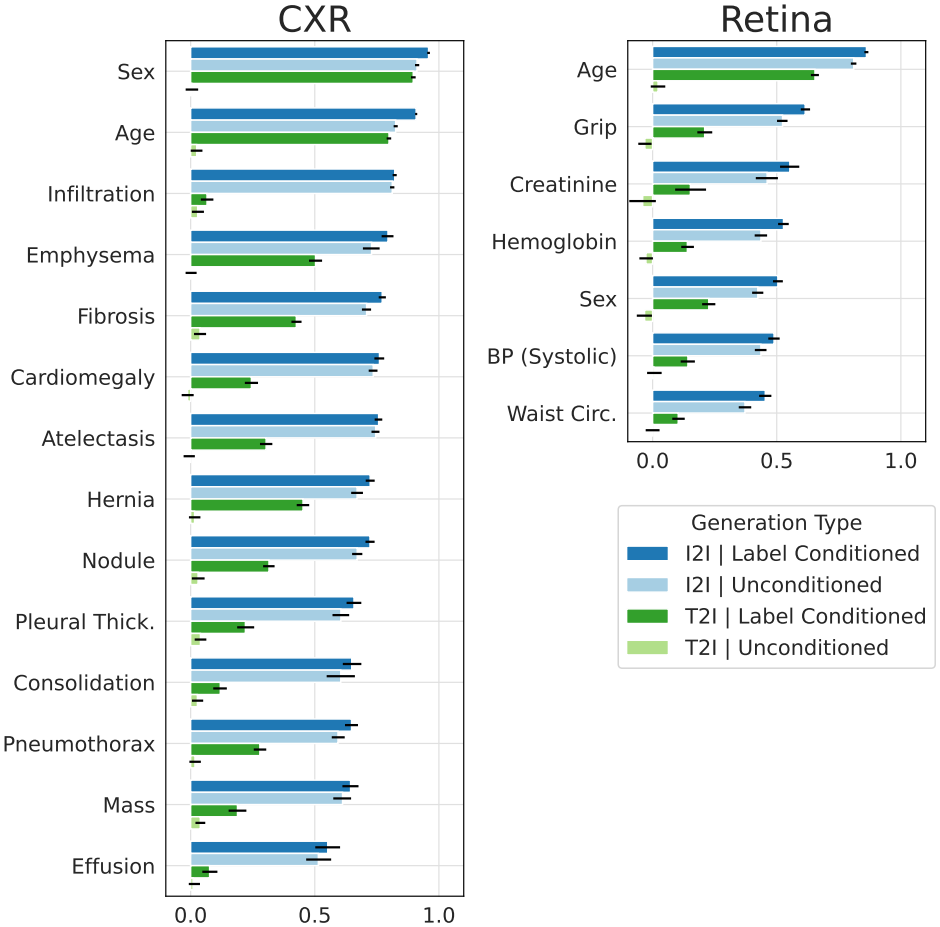
Preservation of clinical signal. Pearson correlation between biomarker estimators on real vs synthetic images for CXR and Retina; error bars, bootstrap 95% CI. Unconditioned, prompt gives image modality only; label-conditioned, prompt includes labels (darker shades; see Section 3.3).

**Figure 3:**
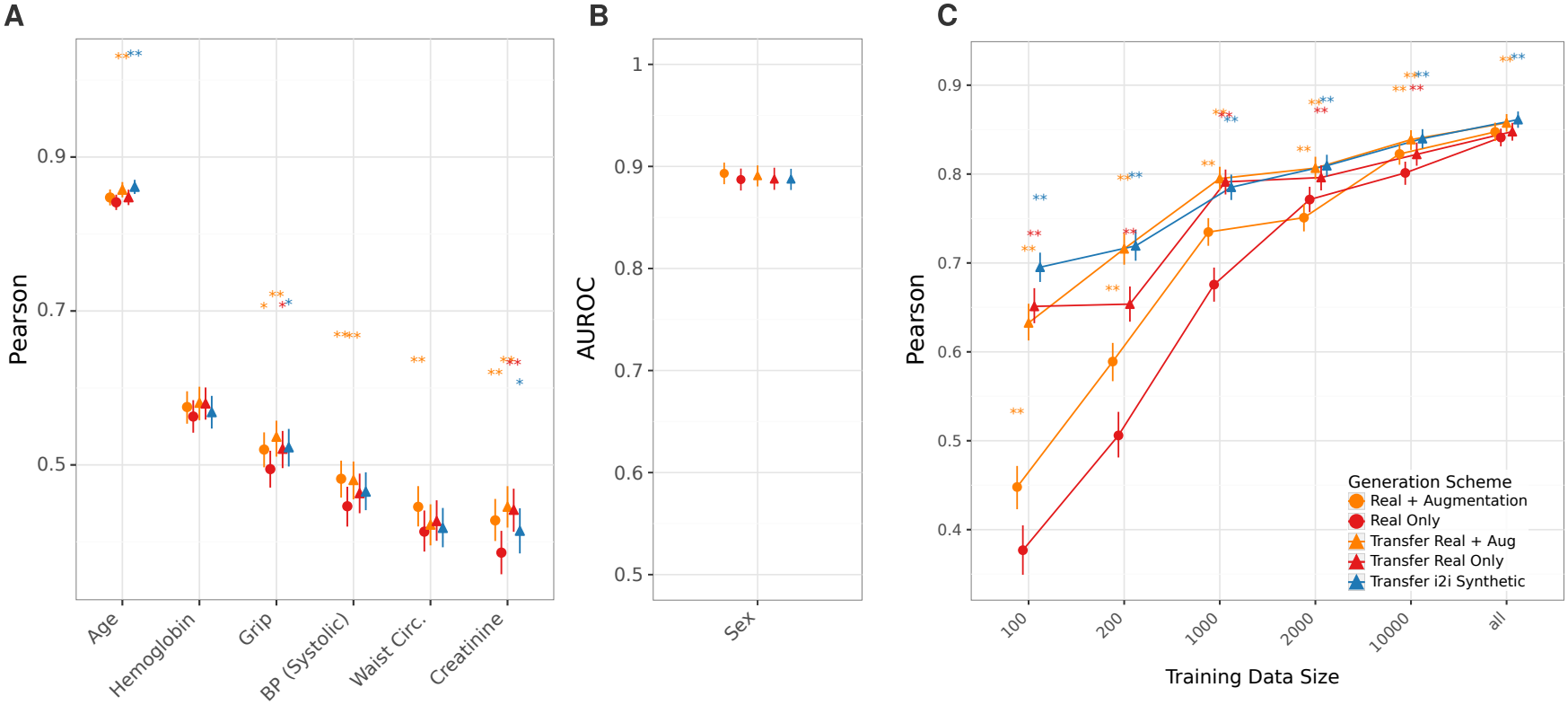
Synthetic-only I2I pretraining transfers across sites at comparable performance to real-image pretraining, with the largest gains in low-data regimes. **a, b**, UKB estimators trained on real (red) or real+augmentation (orange) data, or transfer-learned from I2I-synthetic, real HPP, or augmented real HPP models: **a**, Pearson correlation; **b**, AUROC. **c**, Data-scaling curves for the UKB age estimator under the same schemes. Asterisks: significance vs real-only after FDR correction (0.001 ^***^, 0.01 ^**^, 0.05 ^*^).

**Figure 4:**
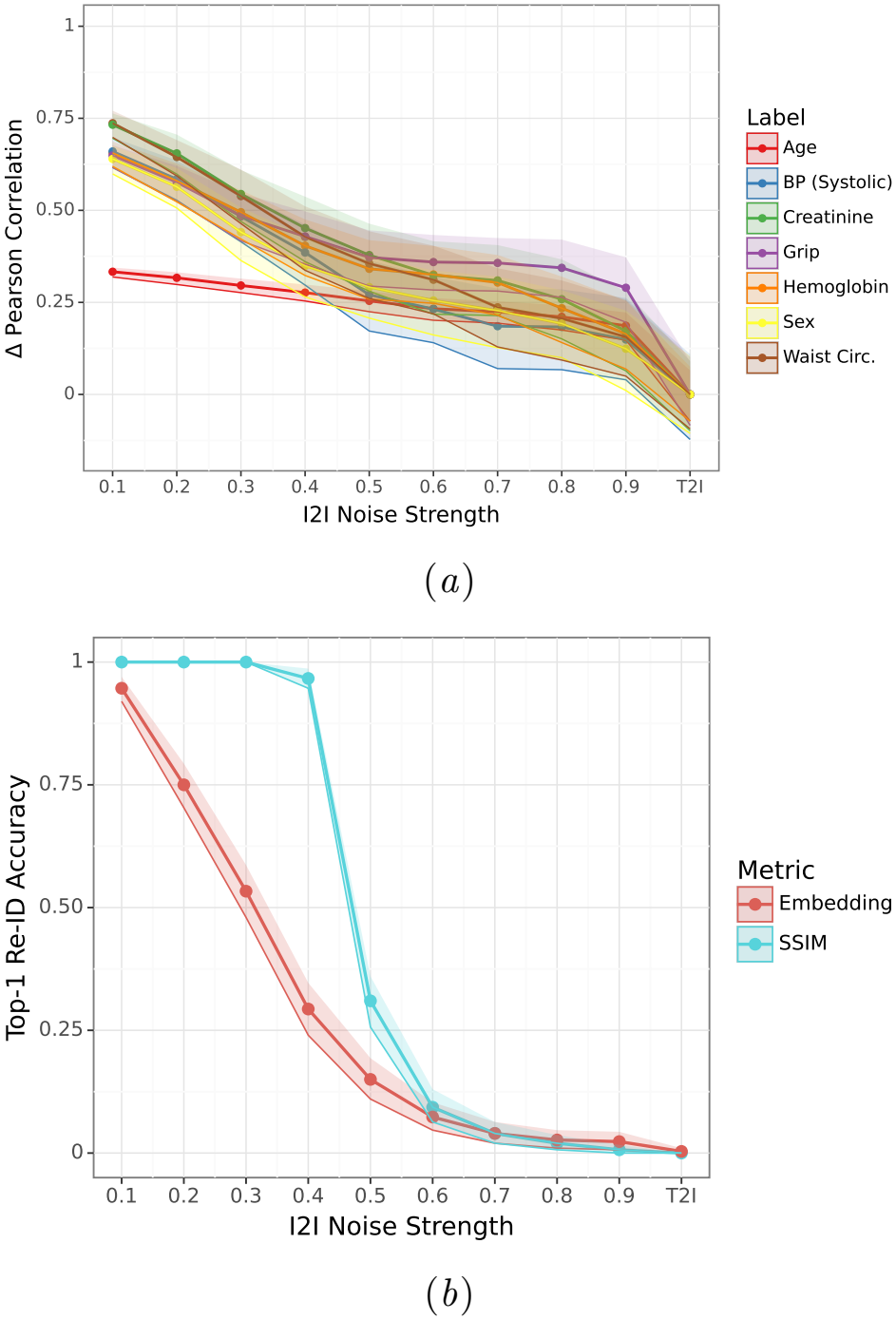
Empirical privacy–utility tradeoff (retina). **a**, Difference between the prediction Pearson correlation (numeric or probability) vs the real image and the same for T2I generation; higher values indicate more original structure retained beyond the prompt. **b**, Risk of identifying the real image by SSIM (blue) or embedding cosine (red) within the test set, vs generation strength (x-axis). Example images across strengths are in Figure 16.

**Figure 5:**
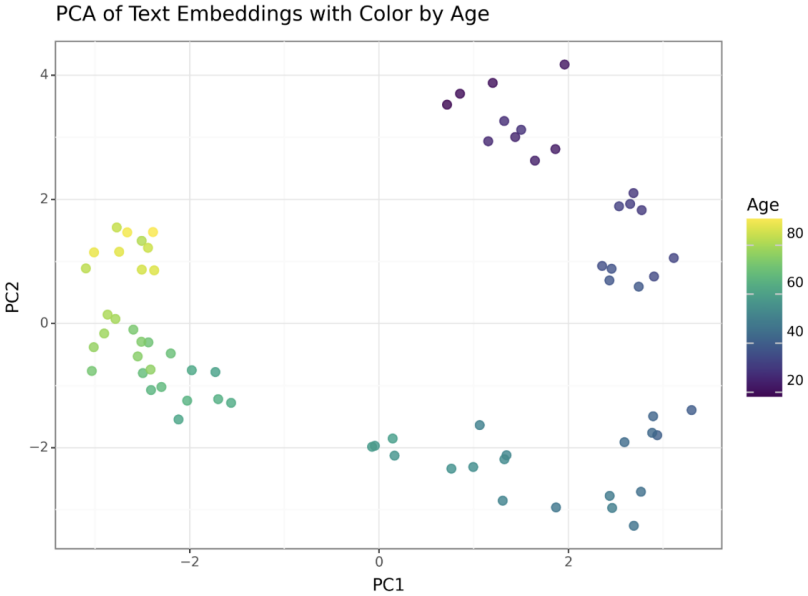
Principal component analysis of prompt text embeddings by age. First (x-axis) and second (y-axis) principal components of prompt embeddings with varying ages 20–90, colored by age. PCA – principal component analysis. See Appendix E.

**Figure 6:**
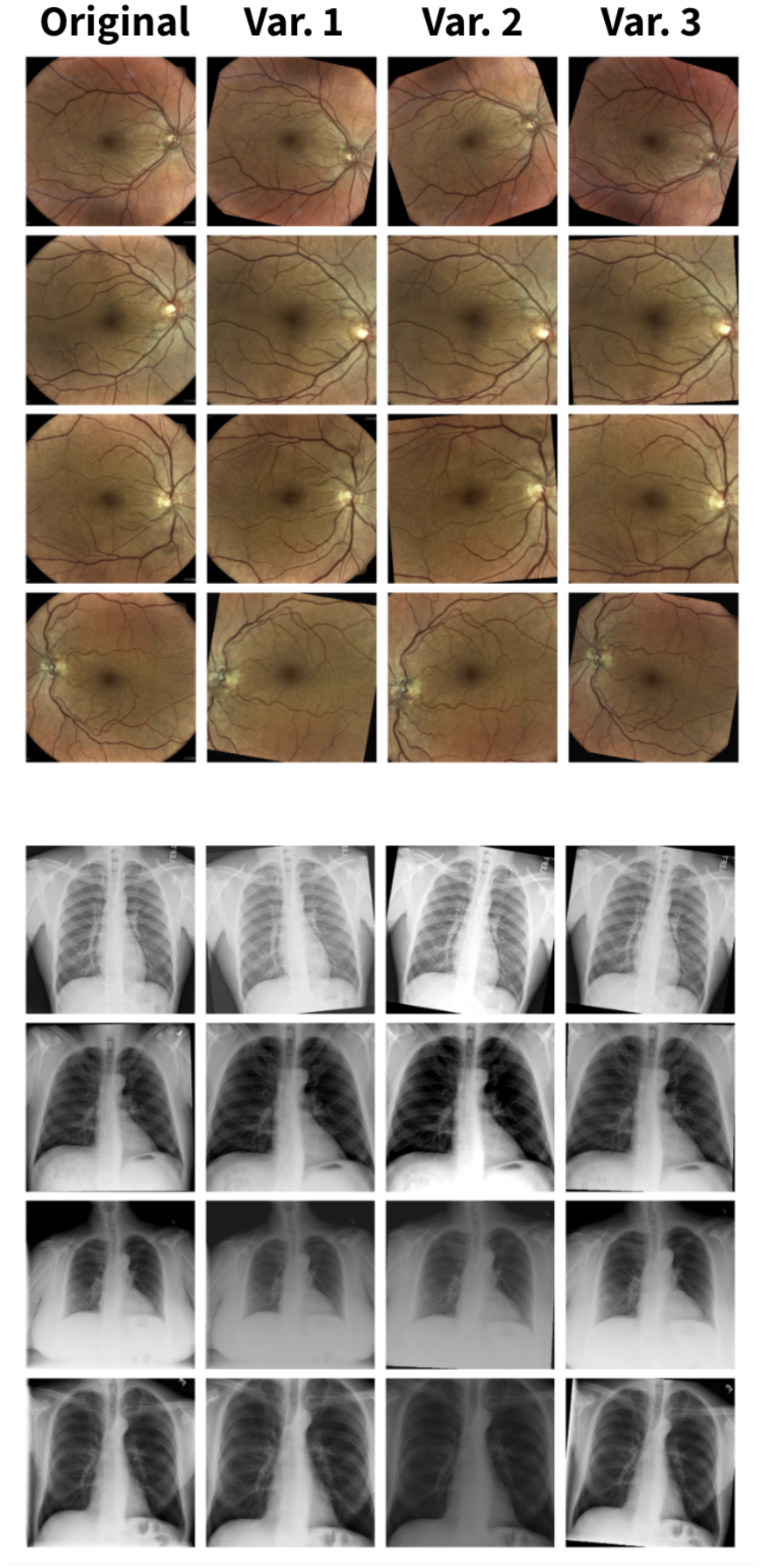
Examples of the conventional augmentation pipeline. Each row corresponds to a single source image: the leftmost column (“Original”) shows the unmodified real image, and columns “Var. 1”– “Var. 3” show three independent stochastic outputs of the same augmentation pipeline (rotation, photometric perturbations, and cropping; see Section B.3). The top four rows show retinal fundus photographs from the Human Phenotype Project; the bottom four rows show chest X-rays from the NIH CXR8 dataset. These augmented variants constitute the conventional-augmentation baseline (orange in Figure 12 and Figure 3) against which synthetic-only and real:synthetic mixed training regimes are compared, and illustrate the visual range introduced by standard augmentations relative to the I2I- and T2I-generated counterparts shown in earlier figures.

**Figure 7:**
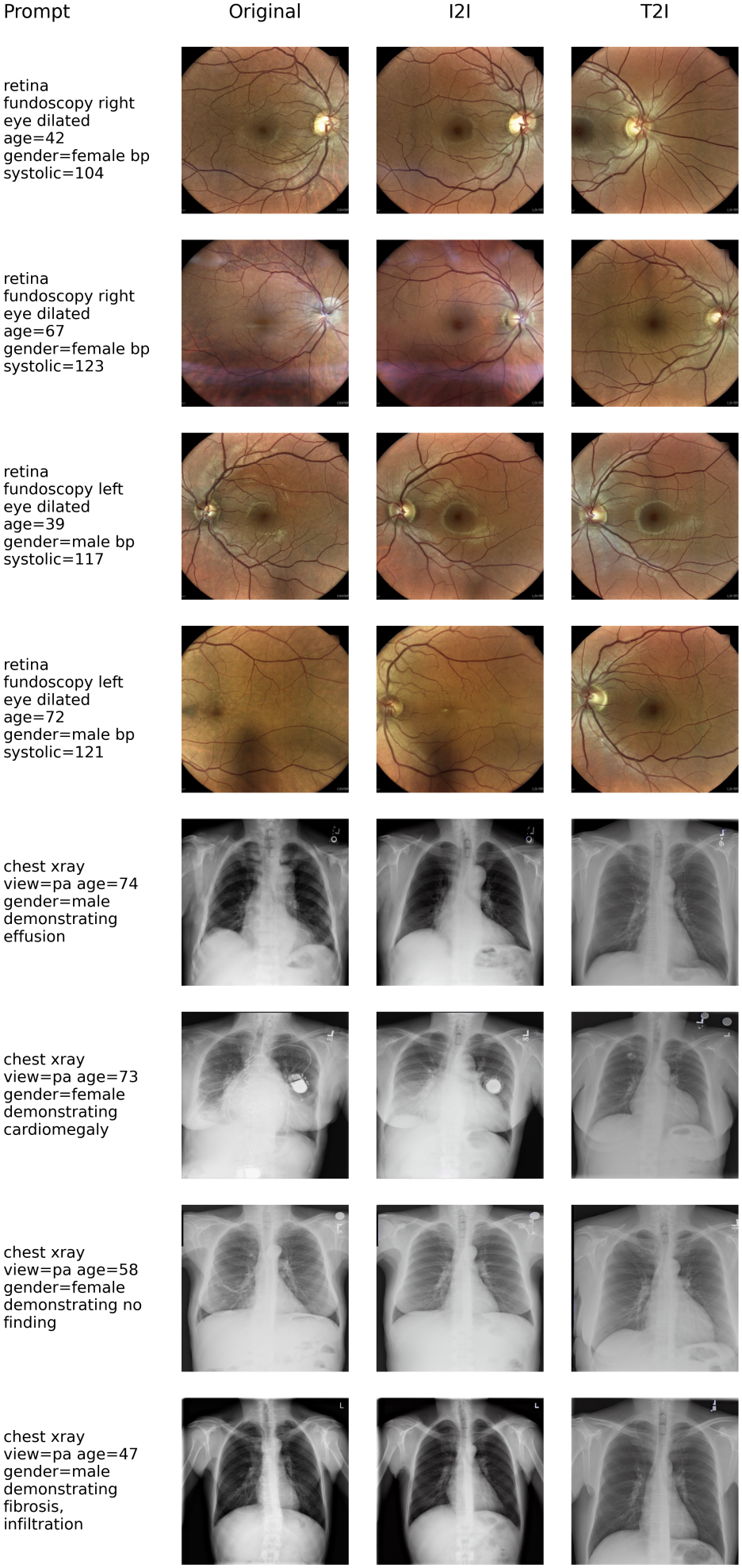
Representative synthesis examples. Retinal fundus photographs (top 4 rows) and chest Xrays (bottom 4 rows). Columns: conditioning text prompt, original real image, image-to-image (I2I) synthetic counterpart, and text-to-image (T2I) synthetic image (same prompt). I2I retains source-specific anatomy (vessel geometry, disc/macula, devices, color) lost in T2I.

To assess whether improved fidelity translates into preservation of clinically relevant information, we compared the agreement of biomarker estimators between real images and their synthetic counterparts. Estimator outputs on I2I images were more strongly correlated with outputs on the original images than those on T2I images across multiple attributes (Figure 2 for retina and CXR images). Estimator predictions on I2I images also agreed more closely with ground-truth labels than T2I predictions for every retinal attribute (Figure 8), most clearly for biomarkers absent from the text-conditioning prompt (hemoglobin, waist circumference, grip strength), indicating that image conditioning retains source-specific information beyond the prompt. This separation is clearest under unconditioned generation, where the prompt specifies only the imaging modality — I2I predictions remained correlated with ground truth whereas T2I collapsed to near zero as expected. The prompted attributes alone, however, predict these biomarkers better than the estimator does on real images (Table 6). After residualizing on them, I2I predictions still tracked real-image predictions, but the estimator itself carried signal beyond these attributes only for hemoglobin and waist circumference (Figure 9 – partial *r* with the true label 0.07 vs 0.19 on real images for hemoglobin; 0.08 vs 0.13 for waist circumference). Finally, when embedding images using DINOv2 embeddings (Section 3.7), the generated I2I images spread across a larger manifold (Figure 10), supporting the interpretation of I2I as a controllable, local transformation.

**Table 6:** Covariate-only baseline for the retinal biomarkers absent from the prompt. Pearson correlation with the true label, mean [95% CI] by bootstrapping. Covariates only: linear regression on the attributes in the label-conditioned prompt (eye, age, sex and systolic blood pressure), fit on the training set. I2I and T2I: estimator on generated images, as in Figure 8. Estimator on real image is predicted on the original image, rest are predicted on generated images. Circ. – circumference.

|  | Covariates | I2I |  | T2I |  | Estimator on |
| Biomarker | only | Label cond. | Uncond. | Label cond. | Uncond. | real image |
|---|---|---|---|---|---|---|
| Hemoglobin | 0.66 [0.64, 0.68] | 0.33 [0.31, 0.36] | 0.21 [0.17, 0.23] | 0.15 [0.12, 0.18] | −0.02 [−0.06, 0.01] | 0.50 [0.46, 0.53] |
| Creatinine | 0.67 [0.63, 0.71] | 0.31 [0.26, 0.36] | 0.16 [0.11, 0.21] | 0.18 [0.11, 0.24] | −0.09 [−0.15, −0.03] | 0.37 [0.31, 0.42] |
| Waist Circ. | 0.52 [0.49, 0.56] | 0.24 [0.21, 0.27] | 0.16 [0.13, 0.18] | 0.07 [0.05, 0.10] | −0.03 [−0.05, −0.01] | 0.35 [0.32, 0.39] |
| Grip | 0.83 [0.81, 0.84] | 0.43 [0.40, 0.45] | 0.27 [0.24, 0.30] | 0.28 [0.24, 0.31] | −0.02 [−0.05, 0.01] | 0.50 [0.47, 0.53] |

**Figure 8:**
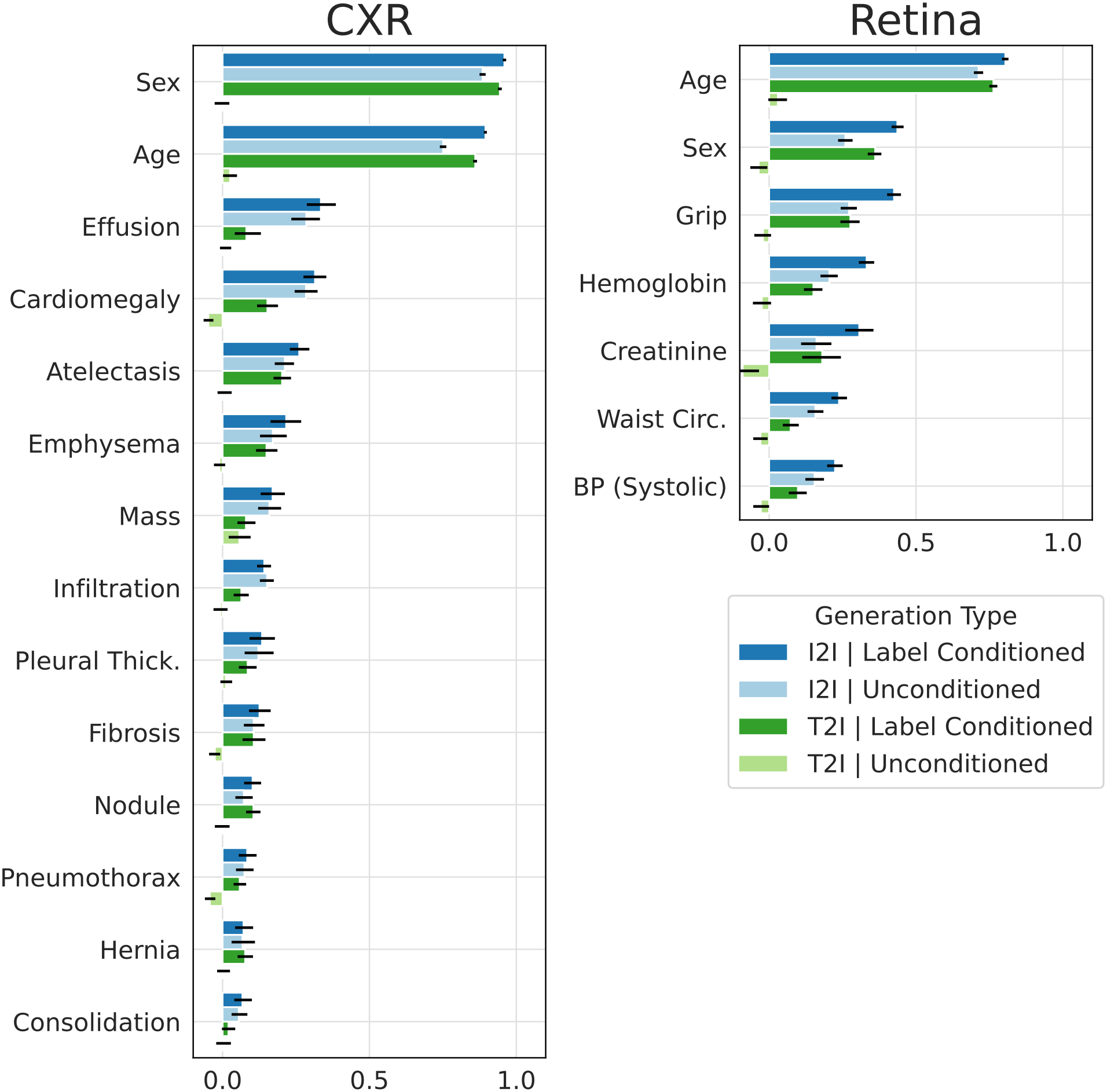
Biomarker estimation from generated images compared to true labels. Pearson correlation of predicted biomarkers on generated images with true labels of the original image. Bars are grouped by generation type – image and text conditioning (blue) and text only conditioning (green); label conditioned — age, sex, etc. (dark blue and green) and unconditioned — containing only modality information (light blue and green). Bars denote 95% confidence intervals by bootstrapping. CXR – chest X-ray; Thick. – thickening; Circ. – circumference; BP – blood pressure.

**Figure 9:**
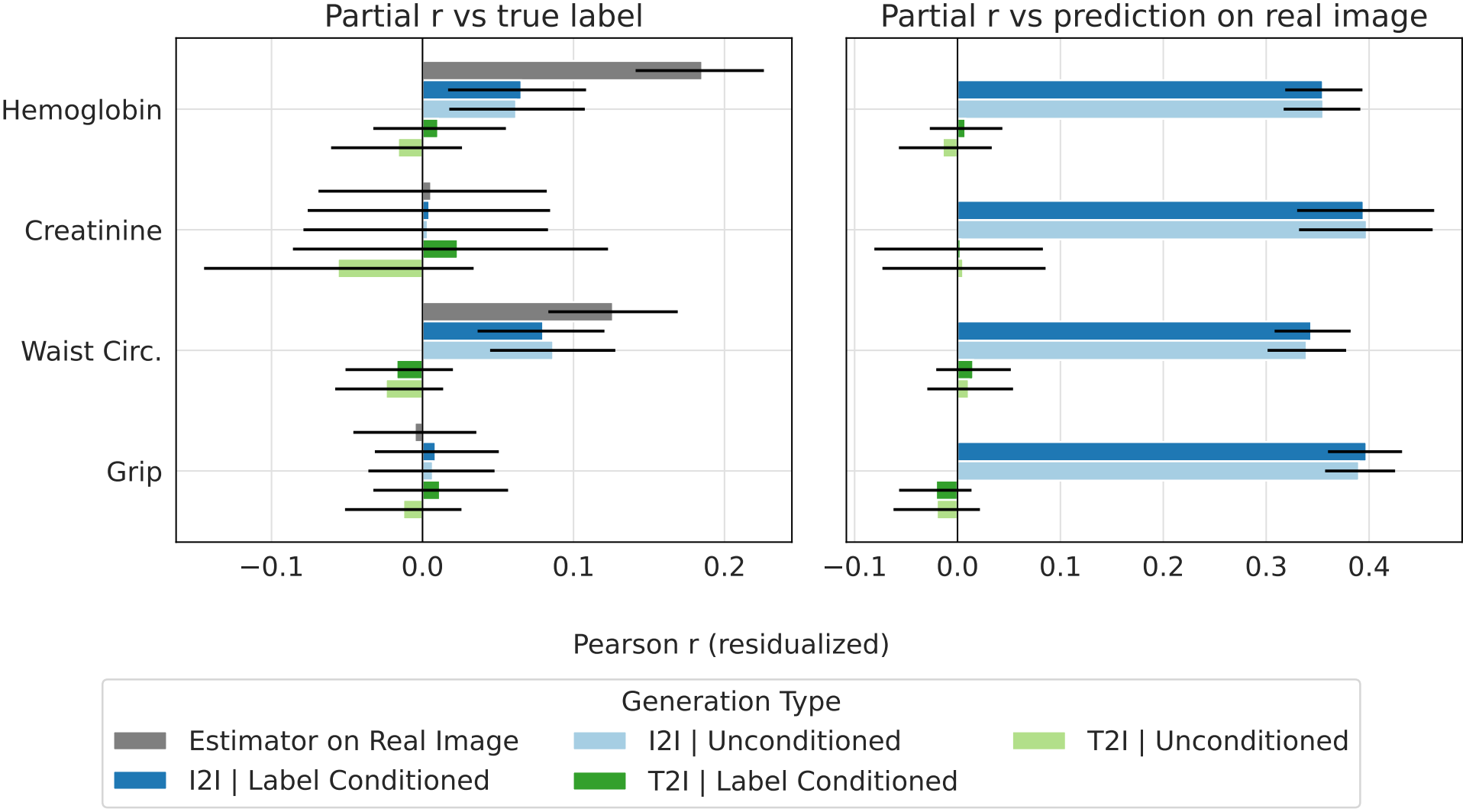
Retinal biomarker signal beyond the prompted attributes. Predictions and true labels were residualized on the attributes in the label-conditioned prompt (eye, age, sex and systolic blood pressure) by linear regression, and Pearson correlation was computed between residuals for the four biomarkers absent from the prompt. **Left**, predictions vs true labels; the estimator on the real image gives the attainable signal. **Right**, predictions on generated images vs predictions on the corresponding real image. Colours as in Figure 8. Bars denote 95% confidence intervals by bootstrapping. Circ. – circumference.

**Figure 10:**
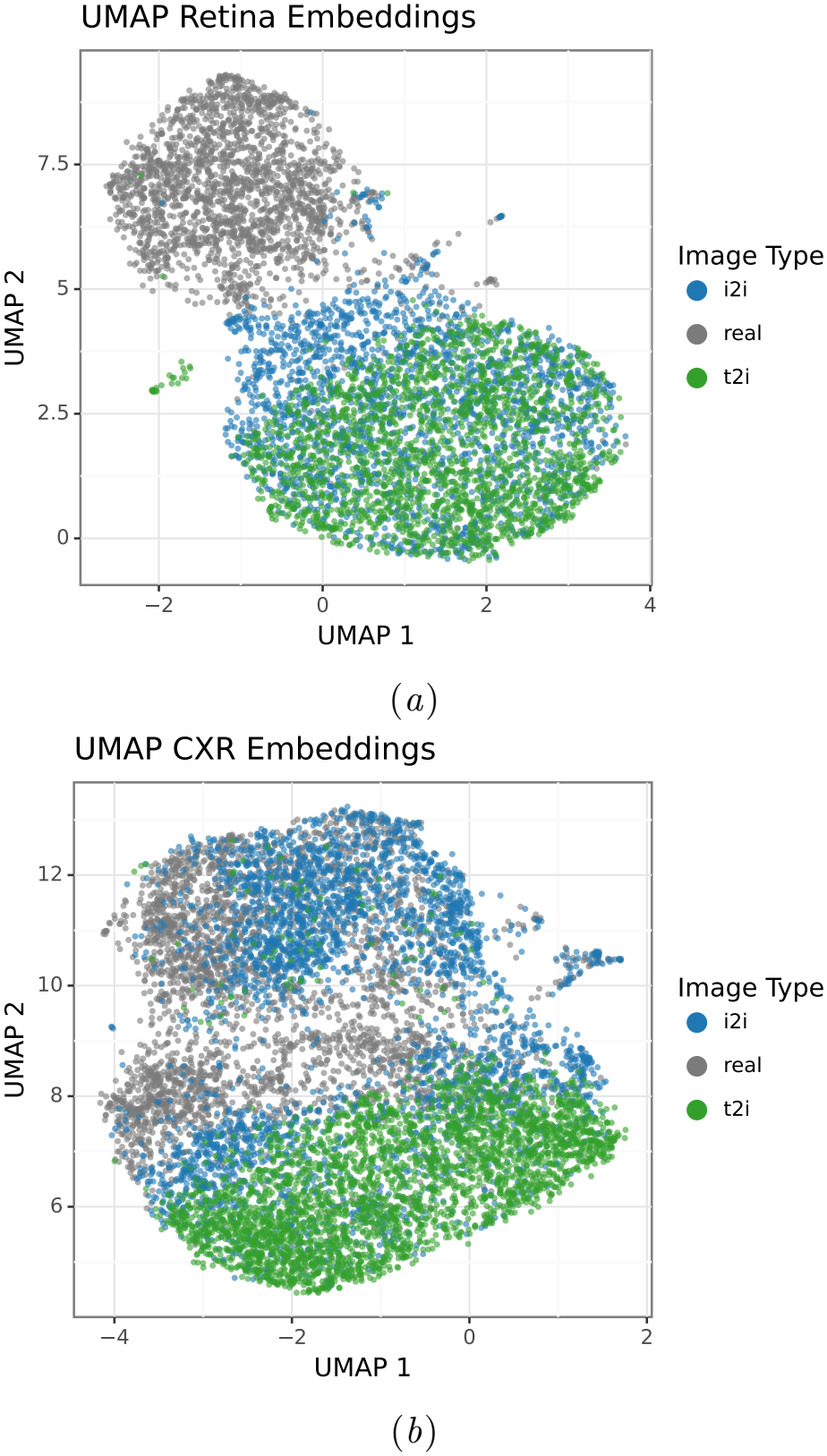
UMAP of self-supervised image embeddings. Two-dimensional UMAP projection of external self-supervised visual features for real images and their I2I and T2I synthetic counterparts, for **a**, Retina and **b**, CXR. I2I generations spread across a larger manifold than T2I, consistent with a controllable, local transformation of each source image.

### 4.2. Training Biomarker Estimators Using Synthetic Images

We next compared estimators trained under the four regimes of Section 3.5 - real-only, real with conventional augmentations, synthetic-only, and mixture of real and synthetic images. While models trained exclusively on synthetic images performed worse than those trained on real images, I2I-derived synthetic training consistently yielded higher predictive performance than T2I-derived synthetic training across the reported attributes (Figure 12A, B; real-data baselines in Figure 11). In 6 of these 7 attributes (all but creatinine), I2I synthetic-only training was sig-nificantly superior to T2I. This suggests that the stronger source-consistency of I2I carries clinically informative variation that is not reliably preserved under text-only generation.

**Figure 11:**
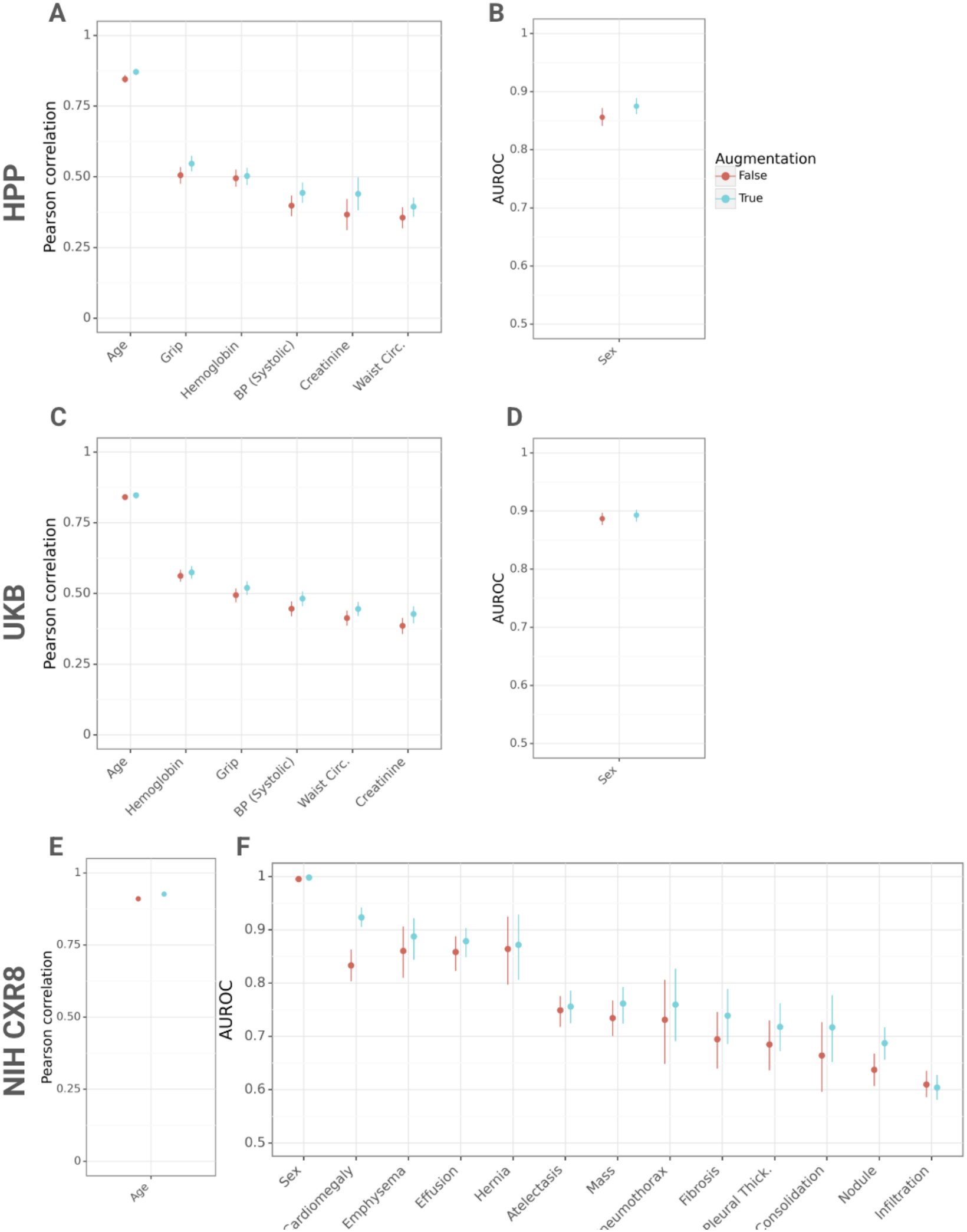
Biomarker estimator model performance, with (blue) and without (red) augmentations, on the test set for models trained on the full training set, measured as Pearson correlation (continuous) or AUROC (binary). **A, B**, HPP retinal fundus biomarkers. **C, D**, UKB retinal images. **E, F**, NIH CXR8 biomarkers and pathological findings. Circ. – circumference; Thick. – thickening.

**Figure 12:**
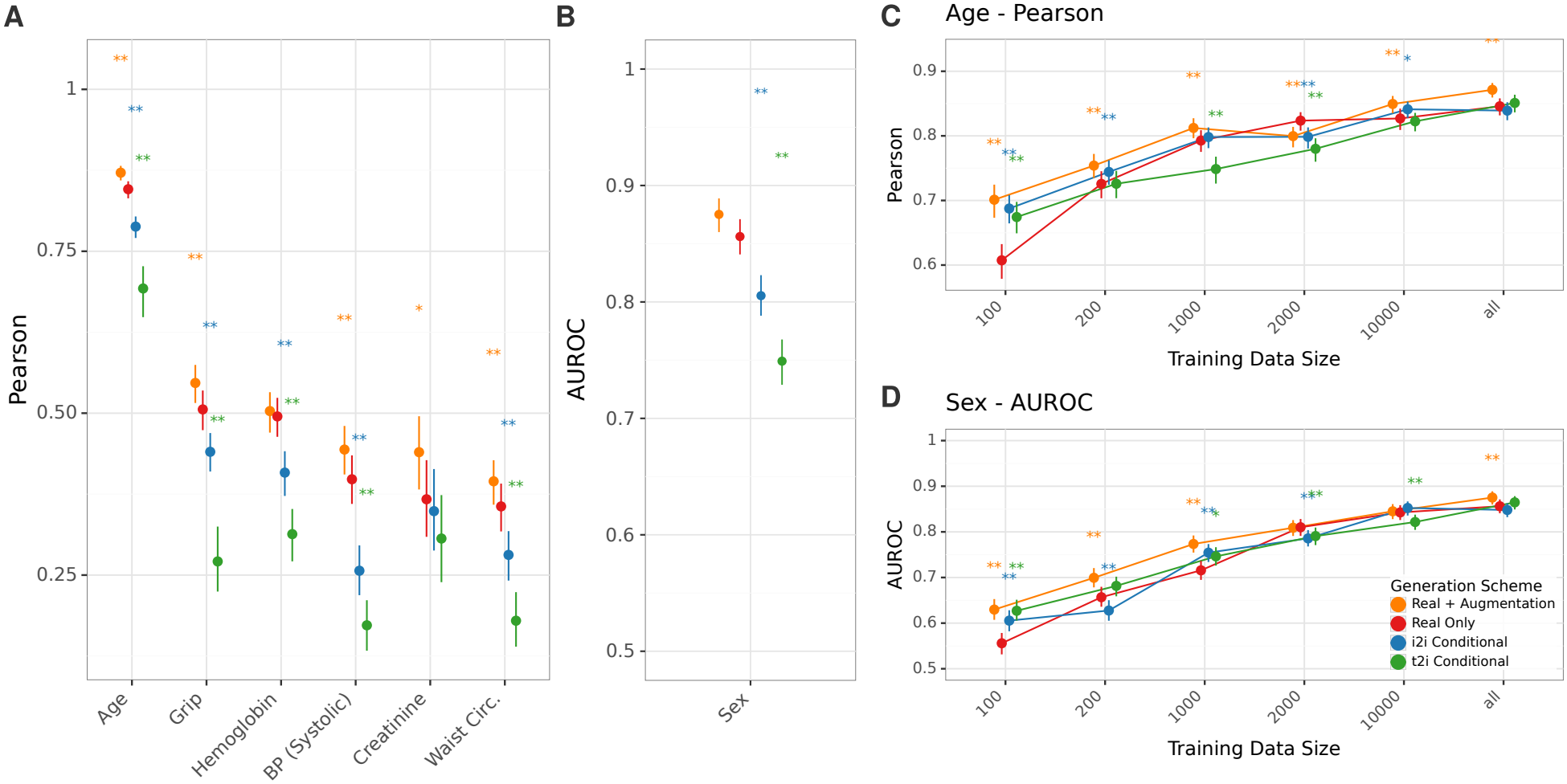
I2I synthetic-only training consistently outperforms T2I across biomarkers. **a, b**, Training on real (red), real+augmentation (orange), I2I-synthetic-only (blue), and T2I-syntheticonly (green) HPP data, conditioned on age, sex, and systolic blood pressure: **a**, Pearson correlation (continuous outcomes); **b**, AUROC (binary outcomes). **c, d**, Training-size scaling curves for age (**c**) and sex (**d**), comparing real-only and real+augmentation baselines with real data mixed with I2I or T2I synthetic images (1:2 real-to-synthetic). Asterisks: significance vs real-only after FDR correction (0.001 ^***^, 0.01 ^**^, 0.05 ^*^).

We further evaluated whether mixing synthetic images with real images improves performance when labeled data are limited. Across subsampled training sizes, adding synthetic images provided the largest relative benefit at small sample sizes, with diminishing gains as the amount of real training data increased. Training with a mix of 1:2 real to synthetic images improved performance when emulating a small training set for both I2I and T2I images in some attributes (age, Figure 12C; sex, Figure 12D; and the rest in Figure 13). We observed no clear trend when training with mixes of 1:1–1:10 real:generated images (Figure 14).

**Figure 13:**
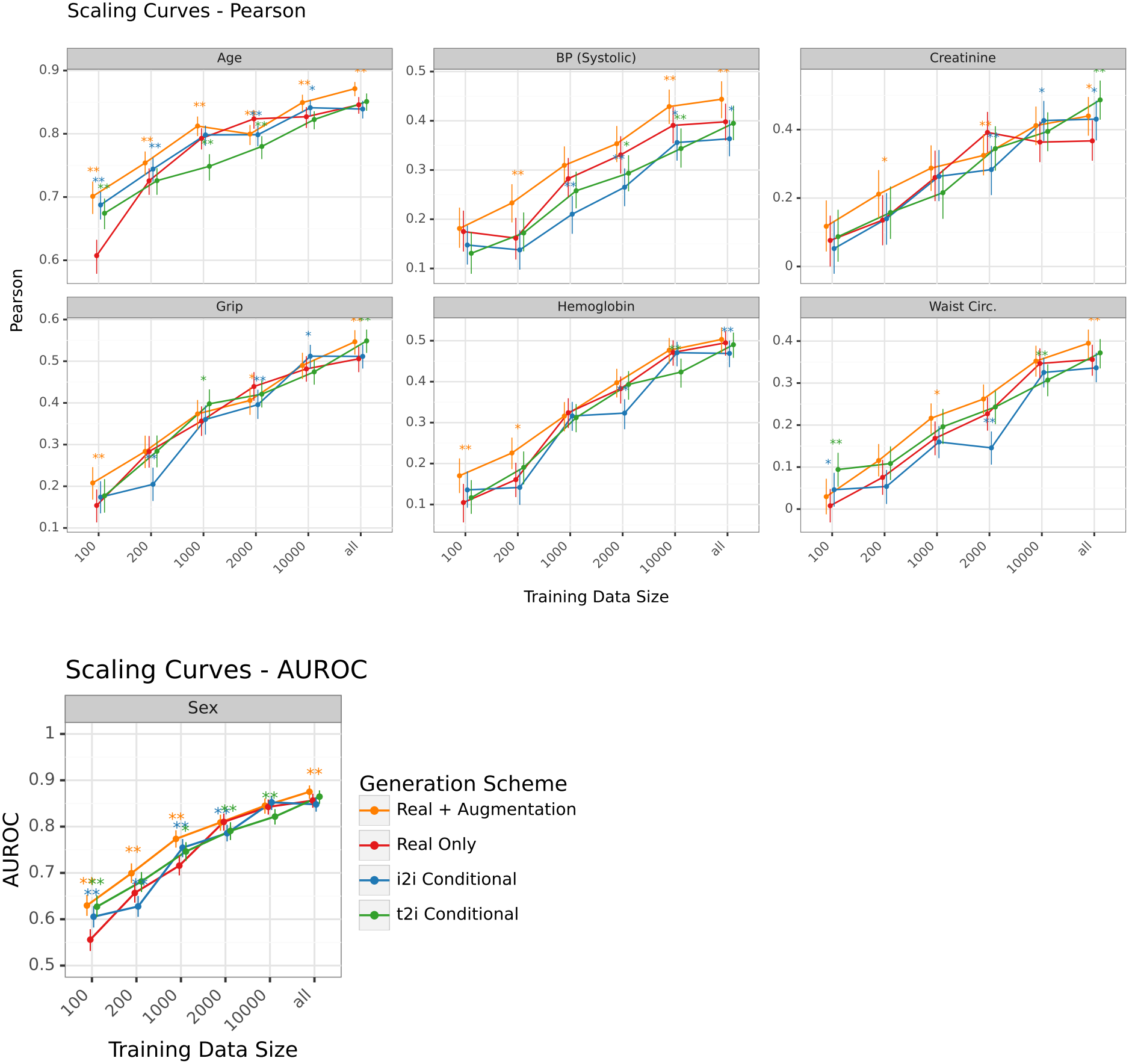
Scaling training with synthetic data. Performance of biomarker estimator models trained on HPP retinal images using only real data (red), real data with standard augmentations (orange), or real images mixed with I2I (blue) or T2I (green) synthetic images at a 1:2 real-to-synthetic ratio across training sizes. Asterisks denote FDR-corrected p-values from bootstrap significance testing compared to real-only training per training size (^*^ *p <* 0.05, ^**^ *p <* 0.01). AUROC – area under the receiver operating characteristic curve; BP – blood pressure; Circ. – circumference.

**Figure 14:**
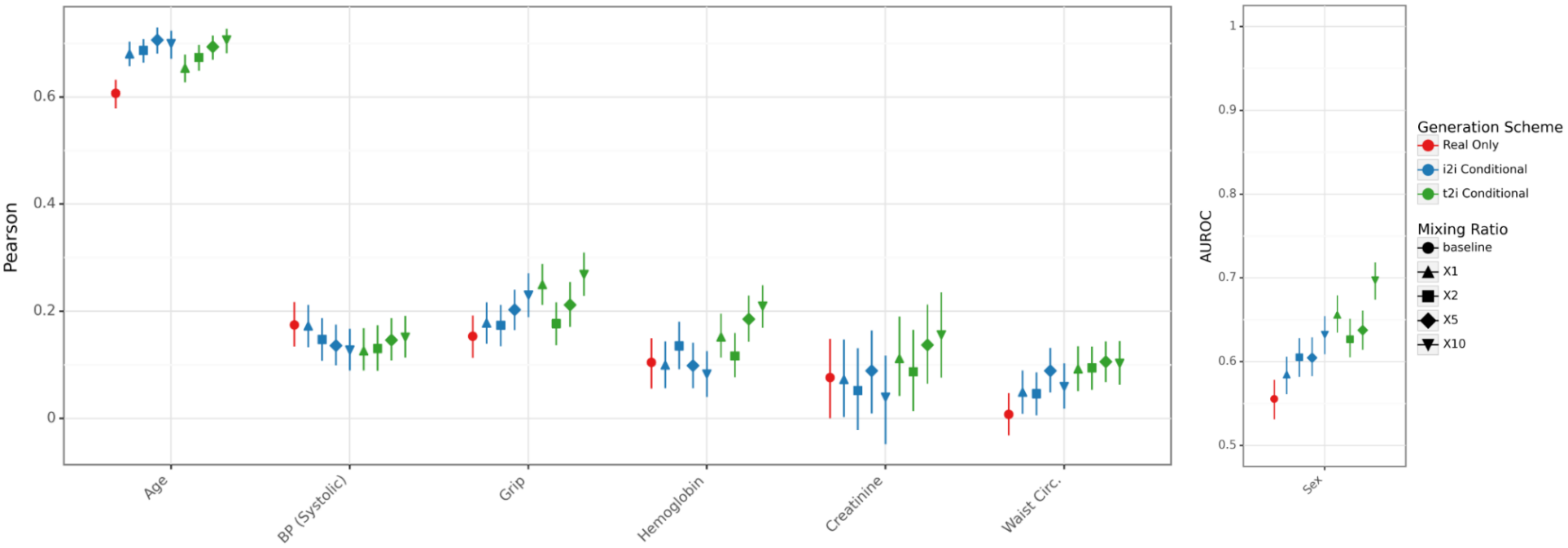
Synthetic data mixing ratio at the low data regime. Performance of biomarker estimator models trained on HPP retinal images using only real data (red), or real images mixed with I2I (blue) or T2I (green) synthetic images at varying real-to-synthetic ratios (1:1–1:10; shape), for training with only 100 real base images. AUROC – area under the receiver operating characteristic curve; BP – blood pressure; Circ. – circumference.

### 4.3. Cross-Site Transfer Learning with Synthetic Data

Cross-site transfer is a major bottleneck in deploying medical image models (Han et al., 2020; DuMont Schütte et al., 2021); we tested whether synthetic images can serve as a substrate for pretraining and transfer, as previously proposed (Liu et al., 2025).

We thus pretrained HPP estimators under each regime and fine-tuned them on UKB (Section 3.5). The two cohorts differed in acquisition conditions and population characteristics (Table 1), providing a clinically relevant domain shift. Models trained on only synthetic I2I images performed comparably to transfer learning from either real or real images with traditional augmentations (Figure 3A–B).

This pattern was stable across UKB fine-tuning set sizes, with the clearest separations observed in the smallest training regimes (Figure 3C for age; Figure 15 for other labels). These findings suggest that synthetic pretraining, particularly with I2I, which preserves source-specific structure, can support crosscohort transfer at a level similar to real-image pretraining, while enabling workflows that reduce dependence on sharing sensitive source images. Notably, the benefit is most clinically relevant in low-data settings where acquiring labeled target-site data is challenging.

**Figure 15:**
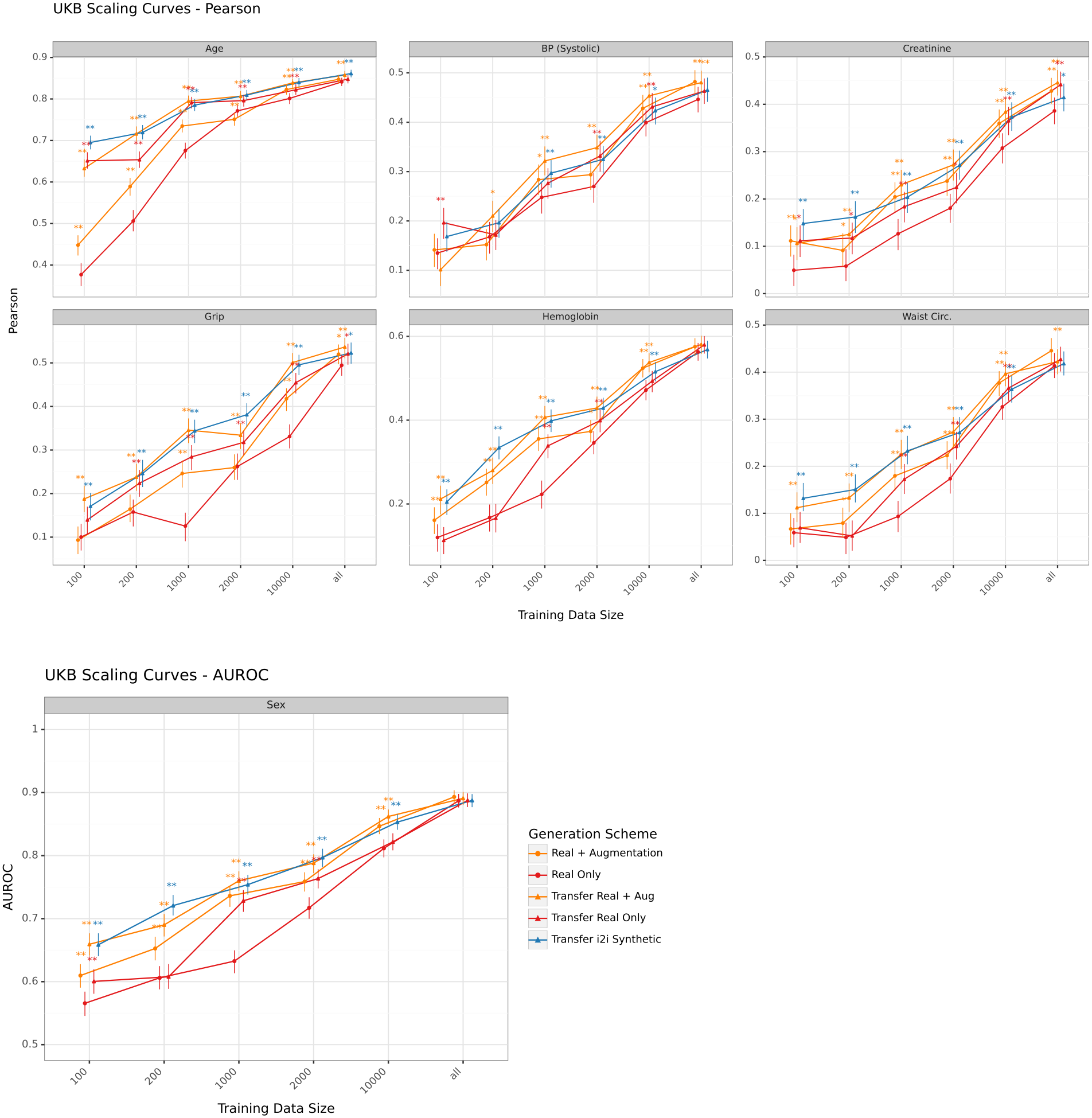
Scaling transfer learning with synthetic data. Performance of biomarker estimator models trained on the HPP retinal training set when transfer-learning to UKB, using only real data (red), real data with standard augmentations (orange), or starting from a model trained (triangles) on real, real + augmentations, or I2I synthetic HPP images; the UKB training phase used real images exclusively. Asterisks denote FDR-corrected p-values from bootstrap significance testing compared to real-only training per training size (^*^ *p <* 0.05, ^**^ *p <* 0.01). AUROC – area under the receiver operating characteristic curve; BP – blood pressure; Circ. – circumference.

### 4.4. Noise Strength Provides a Tunable Control on Re-identification Risk

I2I-generated images show better fidelity but at the potential cost of being more similar to the source image and potentially revealing private information. We therefore quantified a privacy–utility tradeoff that is intrinsic to image-conditioned generation: stronger conditioning preserves more clinically relevant information, but may increase re-identification risk. We sought to quantify the tradeoff between privacy and semantic attribute preservation with respect to the strength *s* of image-conditioning (Section 3.2). In practical terms, increasing strength progressively shifts I2I generation from a near-identity transformation toward a fully synthetic sample as continuous control knob.

As the strength increases, images become more dissimilar to the original (Figure 16) and preserve less of the original attributes (Figure 4A). For example, vessel branching structure and disc features become more dissimilar. Consistent with this, agreement be-tween clinical estimator outputs on the source image and the generated image decreased monotonically with strength across multiple phenotypes. For example, the Pearson correlation for hemoglobin drops from 0.83 at strength 0.1 to 0.52 at 0.5, compared with 0.14 for T2I (Figure 2; Figure 4A). For privacy estimation, we simulated a re-identification attack of 300 synthetic images from the test set against the full test set (Figure 4B). While at strength 0.1, both structural (as measured using SSIM) and semantic distances (as measured using DINOv2 embeddings cosine distance) identify the original image as the nearest image in 100% and 95% of cases, respectively, this drops to 31%, 15% at 0.5, and 2%, 2.7% at 0.8 for SSIM and embedding cosine distances, respectively; true T2I generations from the same prompts link at chance (SSIM 0 and embedding 1 of 300). Scoring recovery at the participant rather than the image level shows similar rates (SSIM 31%, embedding 16.7% at strength 0.5), with similar results for training-split source images matched to the test gallery size (SSIM 27.8%, embedding 9.4%). As a positive control, on real images whose participant was re-imaged in the same eye at a different visit (*n* = 686), the same embedding attack ranked another image of that participant first in 60.8% of cases, confirming the attack’s sensitivity across visits. Notably, the drop in top-1 recovery with increasing noise strength mirrored the decline in attribute preservation, tracing the empirical privacy–utility tradeoff that noise strength makes tunable. Conversely, CXR images (Figure 17) remain highly re-identifiable at the strength used for all utility evaluations: at strength 0.5, SSIM recovers 100% of cases (40.7% by embedding), and remaining high with 63% of cases at 0.9, whereas true T2I generations link none of the 300 sources under either attack. CXR thus represents a boundary case where noise strength alone does not sufficiently reduce linkage.

**Figure 16:**
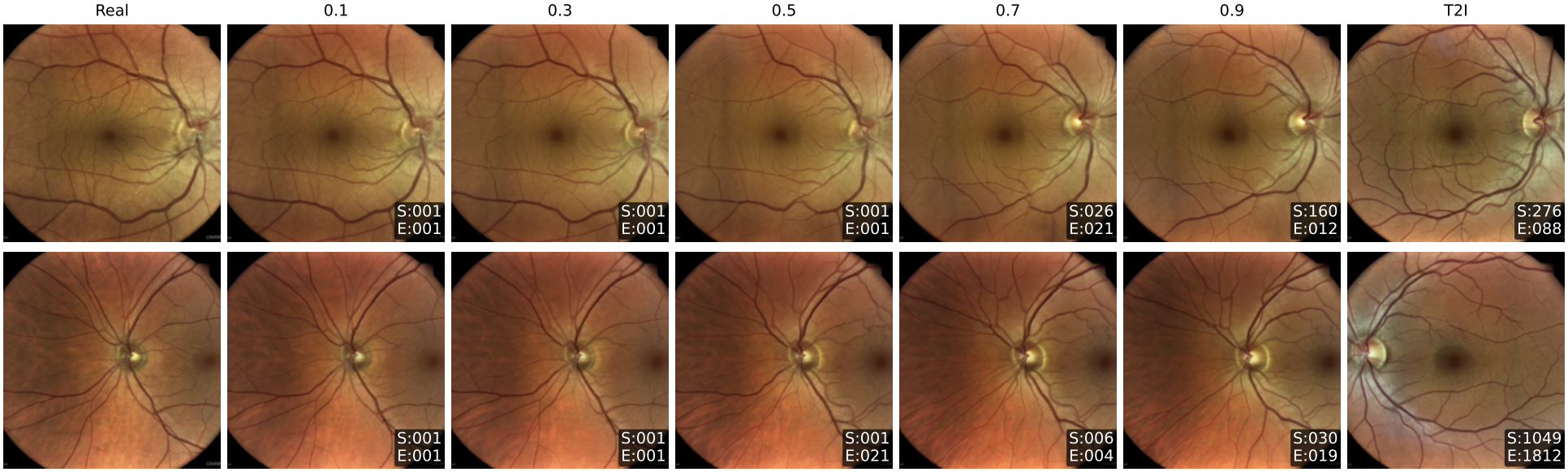
Privacy–utility tradeoff, retina image examples. Two retina examples with I2I generated using a progressive strength schedule from 0.1–0.9, and T2I generation using the same prompt. Insets denote the rank of similarity to the original image in SSIM (S) or embedding cosine similarity (E).

**Figure 17:**
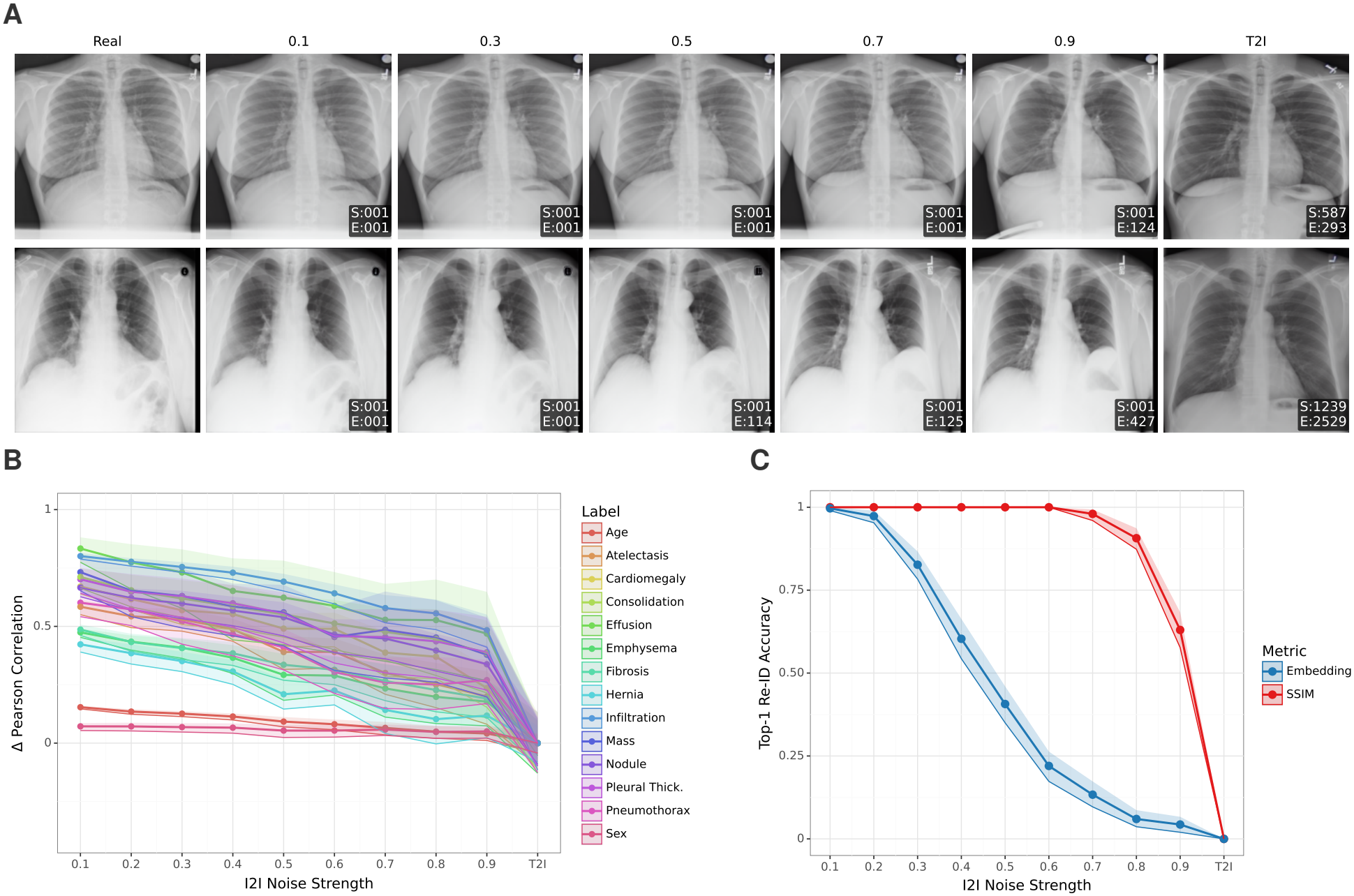
Privacy-utility tradeoff, chest X-ray. **a**, Two example chest X-ray images with I2I generated using a progressive strength schedule from 0.1–0.9, and T2I generation using the same prompt. Insets denote the rank of similarity to the original image in SSIM (S) or embedding cosine similarity (E). **b**, difference between Pearson correlation of predictions (numeric or probability) with the same model predictions on the real image and a similar correlation calculated on the T2I image; a higher value corresponds to higher retained image information than captured in the prompt alone. **c**, Risk of identifying the real image by SSIM (red) or embedding cosine similarity (blue) among the test set, from the generated image at that strength level (x-axis).

The attacks above rely on image similarity alone, whereas a synthetic image is released together with its labels. We therefore also evaluated the attributeaware attackers (Section 3.8; Table 7; same 300 sources and gallery). Matching on these attributes without any image recovers the source in 77% of retinal and 37% of CXR cases, since many of these sources have a unique attribute combination in the gallery (63% and 25%, respectively). This linkage comes from the labels and is the same for every generation method: for T2I, adding the image by either attack leaves recovery at the attribute-only level (em-bedding 78%, 34%), whereas for I2I at strength 0.5 the image raises it to 92% in both modalities (99% and 100% by SSIM). Noise strength thus controls the linkage contributed by the image, while the linkage carried by the released labels calls for separate disclosure control.

**Table 7:** Re-identification with the released labels. Exact-image top-1 (%) on the 300 test sources of the strength sweep (Figure 4B), I2I at strength 0.5 and true T2I generated from the same sources’ prompts. Embedding only: cosine nearest neighbour in DINOv2 embedding space. SSIM only: SSIM nearest neighbour. Attributes only: uniform choice among gallery records whose attributes match the prompt (expected top-1 = mean inverse match-set size). Attributes + embedding (+ SSIM): cosine (SSIM) nearest neighbour among those records. Attributes are eye, age, sex and systolic blood pressure (retina) and view, age, sex and findings (CXR). Gallery: test images, or training and test images (19,075 retinal, 16,222 CXR; no SSIM rows, as the training images’ pixels were not available for this analysis).

|  |  | Retina |  | CXR |  |
|---|---|---|---|---|---|
| Attacker | Gallery | I2I 0.5 | T2I | I2I 0.5 | T2I |
| Embedding only | Test | 15.0 | 0.3 | 40.7 | 0.0 |
| SSIM only | Test | 31.0 | 0.0 | 100.0 | 0.0 |
| Attributes only | Test | 76.9 | 76.9 | 37.3 | 37.3 |
| Attributes + embedding | Test | 92.3 | 77.7 | 92.3 | 34.3 |
| Attributes + SSIM | Test | 99.3 | 75.7 | 100.0 | 35.7 |
| Embedding only | Train + test | 6.0 | 0.3 | 25.0 | 0.0 |
| Attributes only | Train + test | 34.9 | 34.9 | 20.4 | 20.4 |
| Attributes + embedding | Train + test | 73.3 | 38.0 | 82.7 | 19.0 |

A distance to closest record analysis reveals that the distribution of distances between synthetic images and their real counterparts is similar to the distribution of distances between real images for strengths 0.5 and higher for retinal images (Figure 18). At these strengths, synthetic-to-real similarity becomes comparable to baseline real-to-real similarity within the cohort. We emphasize that these evaluations provide empirical estimates of reidentification risk under specific similarity-based attacks, rather than formal privacy guarantees.

**Figure 18:**
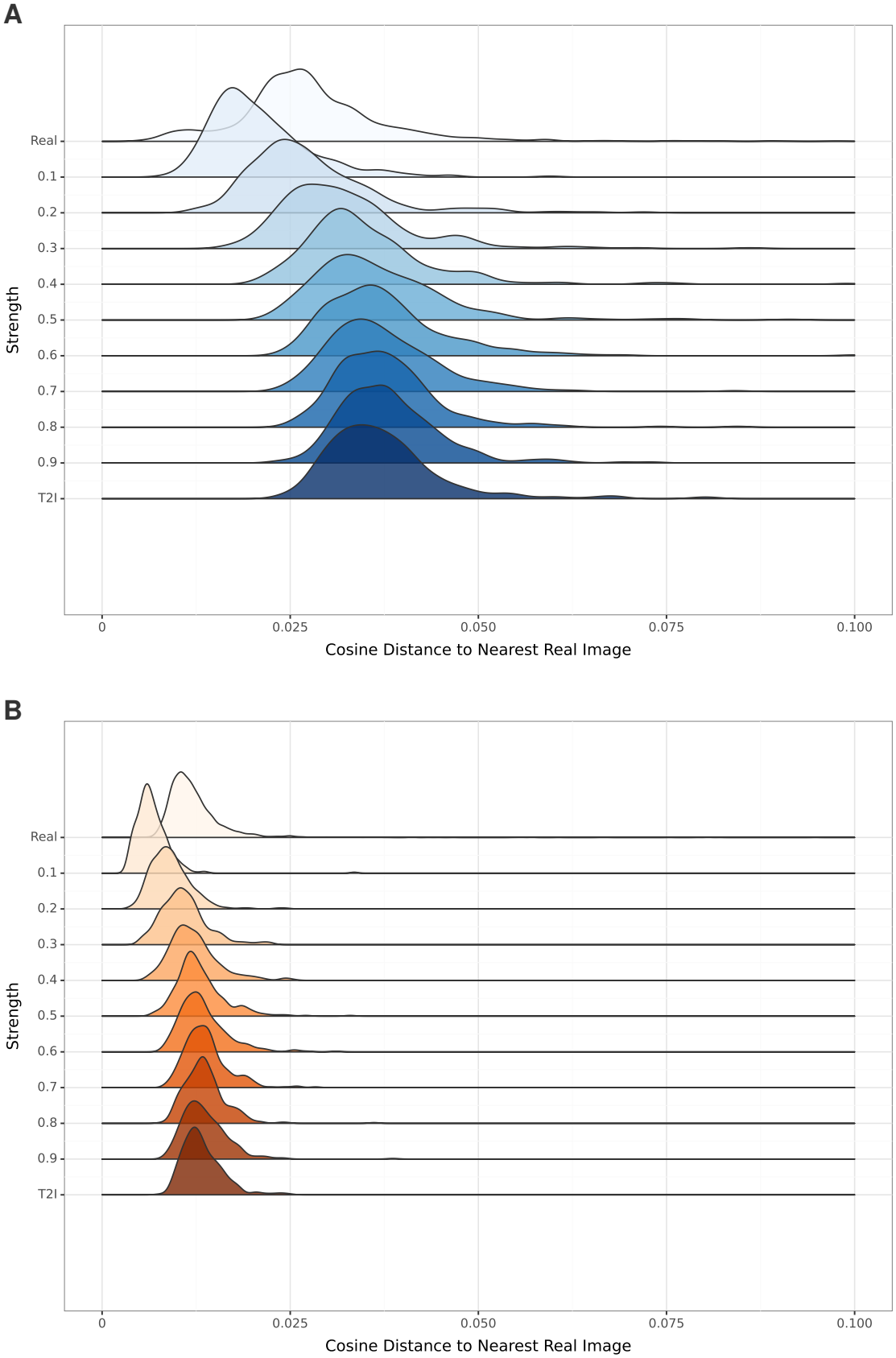
Distance-to-closest-record distributions across I2I noise strengths. **a**, Retina (HPP). **b**, Chest X- ray (NIH CXR8). For each image we computed the cosine distance to its nearest neighbor among real test images using a self-supervised DINOv2 encoder, visualized as a ridge plot stratified by I2I noise strength (0.1–0.9) and for T2I generation. The top “Real” row shows the within-cohort real-to-real DCR baseline (each real test image to its nearest other real image), providing a reference for how close unrelated real images of the same cohort sit in embedding space. For retina (a), synthetic-to-real DCR distributions are broad with long right tails at low noise strength and progressively tighten and shift toward larger distances as noise strength increases; from strength ≈ 0.5 onward, distributions occupy a similar range to the real-to-real baseline rather than collapsing to near-zero. For CXR (b), synthetic-to-real DCR distributions remain tightly concentrated at small distances at every strength, including T2I, overlapping the real baseline throughout. Because T2I generations, which link none of their sources, sit as close as I2I, DCR does not separate the two for chest X-rays; CXR re-identification risk is therefore assessed with the linkage attacks (Figure 17).

## 5. Discussion

We describe image-conditioned diffusion (I2I) for retaining source-specific signal in synthetic medical images with a tunable re-identification risk. On retinal fundoscopy, I2I produces high-fidelity images that re-tain more source-specific signal than T2I—both with and without text-conditioning on the labels—and estimators trained on purely synthetic I2I images outperform those trained on T2I images, though they remain below training on real images. Critically, these synthetic images also support cross-site transfer learning comparable to training on real source-site images. The strength of noise applied to the conditioned image yields a tunable privacy–utility tradeoff, which we quantify as an empirical re-identification risk estimate.

Image-based conditioning methods are prevalent in computer vision since the introduction of SDEdit (Meng et al., 2022) and subsequent works (PérezGarcía et al., 2024). ction has thus far focused mostly on text-conditioning. As we show in this paper, extending diffusion generation to image-conditioned generation retains subtle source-specific features, so that estimates of unprompted attributes such as hemoglobin track the source labels more closely than under T2I. Similarly, while text-based conditioning on continuous attributes is possible and can be represented well (Figure 5), we observed that prompted features, such as age, are also better preserved when the generation is anchored to a specific source image via I2I conditioning, though by a smaller margin.

We present a systematic evaluation of training models utilizing synthetic images, both in-domain and out-of-domain, and with a range of data sizes. Similar to previous works (Sun et al., 2025; Wanget al., 2025; Kim et al., 2024), we observe that synthetic data can supplement real images, specifically in the low data regime. However, in this work, unlike previous ones, synthetic data mixing provides no significant added benefit when compared with traditional image augmentations on the same data. This finding suggests that, for institutions that already have access to the underlying real images, diffusionbased augmentation may not universally outperform well-established augmentation pipelines. In contrast, synthetic data can provide an opportunity for data sharing across institutions. We show that crossdomain transfer learning using a model trained exclusively on I2I synthetic images performs comparably with a model trained on real images with traditional augmentations. This synthetic-only pretraining scenario is particularly relevant when source images cannot be shared due to governance restrictions, but synthetic derivatives or model initializations may be permissible. Sharing synthetic images instead of a model trained on real images leaves the recipient free to choose the architecture, the training objective and the labels, and to pool cohorts; its linkage risk can be audited by the releasing institution before release.

We show that I2I synthetic images can have a reduced, tunable re-identification risk with a controllable trade-off for fidelity and privacy risk. Previous works considered even T2I-generated images re-identifiable (Dar et al., 2026). This difference can be explained by a more relaxed definition of re-identification, which considers all images more similar than expected as memorized. While it is clear that even T2I images are sometimes memorized verbatim (Carlini et al., 2023; Somepalli et al., 2023), our results indicate that the re-identification risk, as measured by nearest-neighbor retrieval, is reduced— though not eliminated—for high-strength I2I retinal images: at strength 0.9, top-1 retrieval recovers the true source in 0.7% (SSIM) and 2.3% (embedding) of cases, far below the near-perfect recovery at low strength but still well above the ∼0.04% expected by chance. It is important to emphasize that our privacy analysis focuses on whether a synthetic image can be matched back to its source record under similarity-based retrieval, which does not by itself mean anonymization (Stadler et al., 2022), rather than establishing formal privacy guarantees as provided, for example, by differential privacy frameworks. However, formal differential privacy approaches may require noise levels during training that would degrade the subtle clinical signals this work aims to preserve. For a controlled release (e.g., under a data-use agreement), this estimate can inform data-sharing decisions.

The relevant privacy threat depends on what is released. (a) *Synthetic I2I images*: the main threat is linking a synthetic image to its source record. This is the scenario our attacks evaluate. (b) *An estimator trained on synthetic images*: membership inference is the main threat, although the estimator has seen only synthetic images. (c) *The fine-tuned generator* : memorization and extraction of its training images become central (Carlini et al., 2023). The cross-site transfer use case corresponds to (a) or (b), with synthetic images shared under a data-use agreement rather than publicly. Within such a release, I2I retains signal that T2I loses, and we suggest choosing the release in two steps (attackers in Section 3.8): first apply disclosure control to the released labels so that they alone link few records to their source, then take the smallest strength at which the attributes + image attackers (attributes + SSIM and attributes + embedding) approach their level on true T2I generations, which carry no source anatomy. With coarsened labels, retinal images approach this level only at the high end of the tested strengths, and CXR images at none (Appendix F).

Chest X-ray images map the boundary of this approach: re-identifiability remains high across all I2I strengths up to 0.9, while true T2I generations link none of their sources under either image-only attack, consistent with chest radiographs being a strong deep-learning biometric (Packhäuser et al., 2022). We speculate there are two main contributing factors. First, the combinatorial nature of CXR labels creates patient subsets that are identifiable by their structured data alone. However, this cannot by itself account for the linkage we observe: the attributes alone recover 37% of sources, and the image-only attacks, which do not use them, recover more (Table 7). Second, unlike biomarkers and identifying features in retinal images, which are diffuse throughout the image, many of the labels in chest X-rays correspond to localized findings. This, in turn, makes random noise less effective at masking these image features, even at high noise levels. Local versus diffuse biomarkers can thus be viewed as a boundary of I2I-based reduction of re-identification risk, suggesting in which modalities the method can be useful. For modalities such as chest X-rays, where anatomy is highly identifying, I2I at any strength we tested does not sufficiently reduce linkage. Possible remedies include masking or regenerating identifying regions, differentially private fine-tuning of the generator for T2I release, and auditing with a CXR-specific re-identification network.

This study is limited by its choice of modalities and labels for each modality. First, extensions to other modalities, including volumetric imaging modalities, are required to span the full breadth of medical imaging. Second, our evaluation emphasizes phenotype prediction and re-identification attacks based on image similarity and released attributes, using a single embedding family and a single CXR dataset. Future work should examine additional privacy threats (e.g., membership inference, extraction audits of the fine-tuned generator, learned re-identification models) and clinical endpoints (e.g., downstream diagnostic classification and calibration). We also did not validate the generated images with clinical experts; a blinded reader study would be most informative for visible findings, such as those in chest X-rays. Third, diffusion generation may preserve site-specific acquisition signatures in ways that are not fully captured by the presented metrics, and further auditing across devices and protocols will be important for deployment. Fourth, we used a single base model for image generation (Stable Diffusion 1.4) and estimator models (BiT-ResNet v2 101). For the generative model, as both retinal and CXR images are significantly out of the pretraining distribution of natural images, we expect little qualitative difference with newer models. For the estimator model, while hyperparameter tuning showed improved performance of ResNet over ViT-based models for biomarker estimation on real images, the robustness of the results for synthetic training remains to be verified. Additionally, as healthcare data sharing is subject to regulatory oversight, it remains to be seen how synthetic data is treated by data governance standards.

Image-conditioned diffusion offers a pragmatic path to medical image derivatives that retain clinically relevant information for downstream modeling with a measurable linkage risk. By making the privacy–utility tradeoff explicit and tunable through the noise strength, it supports both low-data learning and cross-cohort transfer where sharing raw images is infeasible. Future work should extend these evaluations to further modalities and clinical tasks and integrate stronger privacy auditing.

## Appendix A. Dataset Details

This section gives a self-contained description of the three imaging cohorts; summary statistics by cohort and train/test split are reported in Table 1.

### Human Phenotype Project (HPP)

The HPP is a large-scale, prospective cohort encompassing deep clinical and molecular phenotyping of 27,916 participants (Reicher et al., 2025; Shilo et al., 2021). We used the retinal fundus images of 10,010 participants together with clinical data recorded up to or at the fundoscopy visit: age at the visit, sex, heart rate, blood pressure, hip and waist circumference, grip strength, and body mass index (BMI), together with comprehensive laboratory blood panels (hemoglobin, platelets, mean corpuscular volume, white blood cell count, glucose, HbA1c%, creatinine, ferritin, albumin, LDL-, HDL-, non-HDL and total cholesterol, and triglycerides). Including first and subsequent fundoscopy visits gave 23,259 images, with 2.3 (0.8) images and 1.2 (0.4) visits per person (mean (SD)).

#### UK Biobank (UKB)

The UKB includes over 95,000 participants with retinal images (Sudlow et al., 2015), accessed under application ID 28784 and downloaded in 08/2022. We included 17,639 participants from the first instance (0), using only the first visit per participant, for a total of 35,334 images. Following the HPP preprocessing, we took the most recent value of each attribute up to the fundoscopy visit, using the shared attribute set age, sex, hemoglobin, blood pressure, grip strength, waist circumference, and creatinine.

#### NIH CXR-8

NIH CXR-8 is a large public dataset of chest X-rays comprising 108,948 frontal-view images of 32,717 unique participants with 14 textderived labels (Atelectasis, Consolidation, Infiltration, Pneumothorax, Edema, Emphysema, Fibrosis, Effusion, Pneumonia, Pleural thickening, Cardiomegaly, Nodule, Mass, and Hernia) (Wang et al., 2017). The dataset was retrieved from the NIHCC box on 25/04/2025. We removed the Edema and Pneumonia labels, kept the first PA-view image per participant, and downsampled images with no findings to 30% of their number as detailed in the methods section.

#### Image Preprocessing

All images were resized to 512 × 512 and normalized to the range [−1, 1] for use in the diffusion models. Categorical attributes were one-hot encoded and continuous attributes were min-max scaled according to the distribution of the HPP training set (applied to both HPP and UKB).

#### Data Access

The HPP retinal fundus images (Reicher et al., 2025; Shilo et al., 2021) contain personal health information and, in compliance with institutional data-governance requirements, are available to bona fide researchers from universities and other research institutions via a secure cloudbased platform (https://humanphenotypeproject.org/); access instructions are obtained by contacting. UK Biobank retinal images (Sudlow et al., 2015) are available to authorized researchers upon application (https://biobank.ndph.ox.ac.uk/); the present study was conducted under UK Biobank application ID 28784. NIH CXR-8 (Wang et al., 2017) is fully public and is available from the NIHCC box (https://nihcc.app.box.com/v/ChestXray-NIHCC).

## Appendix B. Training Details

### B.1. LoRA Fine-Tuning

We fine-tuned Stable Diffusion 1.4 (Rombach et al., 2022) using Low-Rank Adaptation (LoRA) (Hu et al., 2022) applied to the UNet attention projection layers (query/key/value and output projections), training a small set of adapter parameters while keeping the base model weights frozen, at a fixed image resolution of 512 × 512. LoRA rank was set to 64 with *α* = 32. We trained for 20,000 steps using the AdamW optimizer, a constant learning rate of 1e-4, a batch size of 12, mixed fp16 precision, and no gradient accumulation, saving checkpoints and validation generations at regular intervals to monitor fidelity and conditioning adherence. All LoRA runs were executed with the same data-split boundaries as the downstream estimator experiments.

#### B.2. Estimator Hyperparameter Selection

We performed hyperparameter tuning to optimize the performance of clinical biomarker estimator models trained on real images in the HPP cohort, by grid search over 3,000 training steps using 5-fold cross-validation on the training set. We manually chose configurations based on a combined age-estimation Pearson correlation with the target and sex-estimation AUROC. We tuned the base pretrained-model learning rate ([0, 1e-7, 1e-6, 5e-6, 1e-5, 5e-5, 1e-4]); the estimator-head learning rate ([1e-6, 5e-6, 1e-5, 5e-5, 1e-4, 5e-4]); and the base model variant ([vit base patch16 siglip 224, vit base patch16 siglip 512, resnetv2 101×1 bit]). The selected model was resnetv2 101×1 bit, with base/head learning rates of 5e-5/5e-4 for Retina and 1e-4/1e-4 for CXR.

#### B.3. Image Augmentations

We applied image augmentations using torchvision transforms on the fly as part of data loading, with each augmentation applied at a specific rate. We preconfigured and manually tuned the pipelines a priori to achieve medically plausible images that respect the image labels; for example, we refrained from horizontal flipping of retinal images to respect the eye-side label. The resulting pipelines are given in Table 4.

## Appendix C. Image Generation

For each LoRA model, synthetic images were generated with both the T2I and I2I pipelines implemented in the Hugging Face Diffusers framework. Unless otherwise specified, we used the DDIM scheduler and low classifier-free guidance (guidance scale 2) based on generation-parameter optimization sweeps. For I2I generation, we evaluated a range of noise strengths spanning low to high transformation, and for each strength produced multiple stochastic variants per source image by varying the random seed. All generated images were stored with accompanying metadata (prompt, strength, and source image identifier for I2I) to enable full reproducibility and to support linkage analyses; as a basic safeguard against trivial duplicates we performed exact-duplicate checks (e.g., filename hashing). The I2I source images used throughout the fidelity and physiological-signal analyses are drawn from the held-out test split (2,342 retinal images; Table 1), which is disjoint from the LoRA training split, no evaluated source image was seen during generator fine-tuning.

### C.1. Prompt Templates

Text conditioning followed structured prompt templates based on the image modality (retina funduscopy, chest X-rays) and experiment condition (label conditioned or not).

Written as Python format strings pseudo-code, where sex is “female”/”male”, and CXR findings are one of: effusion, cardiomegaly, emphysema, mass, pneumothorax, hernia, atelectasis, pleural thickening, nodule, fibrosis, consolidation, infiltration.

## Appendix D. Computational Resources

All training, synthetic image generation, and evaluation was performed on the Weizmann GPU cluster consisting of NVIDIA L40, A10 and A100 GPUs. We used Python 3.11, CUDA 11.8, and PyTorch 2.7 for all code executed.

## Appendix E. Textual Embeddings of Continuous Measures

It is unclear how coherent continuous measures are embedded through the text encoder of diffusion models. As a heuristic, we embedded prompts with varying ages and projected them using principal component analysis to 2 dimensions. We embedded the prompt “retina fundoscopy right eye dilated age=X gender=male” for all integer values of 20 ≤ *X <* 90 using the CLIP text encoder of Stable Diffusion, resulting in 768-dimensional embedding vectors. The PCA-projected 2D representation found in Figure 5 shows that prompts for close ages are embedded closely in this space.

## Appendix F. Release Criterion with Coarsened Labels

Section 5 proposes choosing an I2I release in two steps. First, apply disclosure control to the released labels so that the labels alone link few synthetic images to their source records. Second, choose the smallest I2I strength at which attackers that combine the released labels with the image approach their recovery rate on true T2I generations, which carry no source anatomy. Here we apply this criterion with the attackers of Section 3.8, on the same 300 test sources and test gallery as Table 7.

For the first step, we coarsened the retinal labels, binning age in 10-year bands and systolic blood pressure in 20 mmHg bands. This lowers attribute-only linkage from 77% with exact labels to 3%. For CXR, we binned age in 10-year bands.

For the second step, we used the attributes + SSIM attacker. This attacker restricts the gallery to records whose released (here, coarsened) attributes match those of the synthetic image, then picks the record whose image has the highest SSIM to the synthetic image. On I2I images, attributes + SSIM recovers more sources than attributes + embedding (99% vs 92% for retina and 100% vs 92% for CXR at strength 0.5 with exact labels; Table 7), so it is the stricter of the two attackers that combine attributes with the image. As the reference level, we took the top-1 recovery of the stronger combined attacker on T2I generations from the same prompts: 5% for retina and 17% for CXR.

For retina, I2I recovery exceeds this reference by +81%, +24% and +10% at strengths 0.5, 0.7 and 0.9, respectively. Retinal images therefore approach the T2I level only at the high end of the tested range, where attribute fidelity is also lower (Figure 4A). For CXR, recovery exceeds the reference by +70% or more at every tested strength (87% vs 17% at strength 0.9), so no tested strength meets the criterion.

The criterion is defined relative to the attackers evaluated here, and it applies to a controlled release (e.g., under a data-use agreement).

